# MSstatsPTM: Statistical relative quantification of post-translational modifications in bottom-up mass spectrometry-based proteomics

**DOI:** 10.1101/2022.09.24.509068

**Authors:** Devon Kohler, Tsung-Heng Tsai, Erik Verschueren, Ting Huang, Trent Hinkle, Lilian Phu, Meena Choi, Olga Vitek

## Abstract

Liquid chromatography coupled with bottom up mass spectrometry (LC-MS/MS)-based proteomics is increasingly used to detect changes in post-translational modifications (PTMs) in samples from different conditions. Analysis of data from such experiments faces numerous statistical challenges. These include the low abundance of modified proteoforms, the small number of observed peptides that span modification sites, and confounding between changes in the abundance of PTM and the overall changes in the protein abundance. Therefore, statistical approaches for detecting differential PTM abundance must integrate all the available information pertaining to a PTM site, and consider all the relevant sources of confounding and variation. In this manuscript we propose such a statistical framework, which is versatile, accurate, and leads to reproducible results. The framework requires an experimental design, which quantifies, for each sample, both peptides with post-translational modifications and peptides from the same proteins with no modification sites. The proposed framework supports both label-free and tandem mass tag (TMT)-based LC-MS/MS acquisitions. The statistical methodology separately summarizes the abundances of peptides with and without the modification sites, by fitting separate linear mixed effects models appropriate for the experimental design. Next, model-based inferences regarding the PTM and the protein-level abundances are combined to account for the confounding between these two sources. Evaluations on computer simulations, a spike-in experiment with known ground truth, and three biological experiments with different organisms, modification types and data acquisition types demonstrate the improved fold change estimation and detection of differential PTM abundance, as compared to currently used approaches. The proposed framework is implemented in the free and open-source R/Bioconductor package *MSstatsPTM*.

## Introduction

Signaling mechanisms allow cells to mount a fast and dynamic response to a multitude of biomolecular events. Signaling is facilitated by the modification of proteins at specific residues, acting as molecular on/off switches [12, 10, 3]. Characterizing relative abundance of a modification site’s occupancy repertoire across experimental conditions provides important insights [25]. For example, meaningful patterns of changes in post-translational modifications (PTMs) abundance can serve as biomarkers of a disease [30]. Alternatively, distinguishing the quantitative changes in a PTM from the overall changes of the protein abundance helps gain insight into biological and physiological processes operating on a very short timescale [40, 6, 20]. This helps to distinguish between relative site occupancy changes at steady-state protein levels, typical for short time-scale signaling events, and observed relative changes of PTMs as a result of underlying gene expression or protein abundance levels.

Bottom-up liquid chromatography coupled with tandem mass spectrometry (LC-MS/MS) is a tool of choice for unbiased and large-scale identification and quantification of proteins and their PTMs [17, 32]. However, LC-MS-based interrogation of the modified proteome is challenging, for a number of reasons. First, the relatively lower abundance of modified proteoforms dictates that a global interrogation can only be achieved through large-scale enrichment protocols with modification-specific antibodies or beads [15]. Variability in the enrichment efficiency inevitably affects the reproducibility of the number of spectral features (e.g., peptide precursor ions or their fragments) and their intensities. Second, contrary to the often large number of identified peptides that can be used to quantify protein abundance, there are relatively few representative peptides that span a modification site, and there may be multiple modified sites on a single peptide [25]. Third, unless early signaling events are interrogated, the interpretation of the relative changes in modification occupancy are inherently confounded with changes in the overall protein abundance, complicating the interpretation of the results [40, 29]. Finally, technological aspects of bottom-up MS experiments, such as presence of labeling by tandem mass tag (TMT), introduce additional sources of uncertainty and variation.

The technological difficulties in PTM identification and quantification increase the uncertainty and the variation in the data, and challenge the downstream statistical analyses. Frequently data from these experiments are analyzed using statistical methods that were not originally designed for this task. Researchers use methods such as *t*-test[18], Analysis of Variance (*ANOV A*)[14], or *Limma*[31], by taking as input the intensity ratios of modified and unmodified peptide summaries, and comparing the mean abundance of different PTM sites. Such approaches do not fully account for all the sources of uncertainty. As the result, these approaches are either not directly applicable to experiments with non-trivial designs (such as experiments with multiple conditions, paired and time course designs, and experiments with labeling), or require the analysts to exercise non-trivial statistical expertise.

This manuscript proposes a versatile statistical analysis framework that accurately detects relative changes in post-translational modifications. The framework requires an experimental design, which quantifies, for each sample, both the peptides with post-translational modifications, and peptides from the same proteins with no modification sites. The framework supports data-dependent acquisitions (DDA) that are label-free or tandem mass tag (TMT)-based. The statistical methodology separately summarizes the abundances of peptides with and without the modification sites, and fits separate linear mixed effects models that reflect the biological and technological aspects of the experimental design. Unmodified peptides may or may not span a modifiable site. Next, model-based inferences regarding the PTM and the unmodified protein-level summaries are combined to account for the confounding between these two sources.

We evaluated the proposed framework on two datasets from computer simulations, one benchmark controlled mixture, and three biological investigations. The datasets illustrate a diverse set of organisms, modification types, acquisition methods and experimental designs, showing the applicability of the framework to a variety of situations. By appropriately leveraging the information from the unmodified portion of the protein sequence, the proposed approach improved the accuracy of the estimates of PTM fold changes, and produced a better calibrated false positive rate of detecting differentially abundant PTMs as compared to existing methods. In particular, accounting for the confounding from unmodified protein abundance allowed us to characterize the true effect of the modification, avoiding the need for more manual and time intensive follow-up investigation.

The proposed approach is implemented as a freely available open source R package *MSstatsPTM*, as part of the *MSstats* family of packages [9, 16], and is available on Bioconductor.

## Experimental procedures

### Data overview and availability

Table 1 summarizes the experiments. Two computer simulations had known ground truth, and varied in experimental realism. The first simulation produced a perfectly clean dataset, with many replicates and no missing values. The second simulation introduced real-world characteristics, such as limited modified features and missing values. Details of computer simulations are available in **Supplementary Sec. 2.1 & 2.2**, and on GitHub (https://github.com/devonjkohler/MSstatsPTM_simulations).

**Table 1:** Simulated and experimental datasets. “Dataset” is the dataset code name. “No. of bio. replicates” shows the number of biological replicates per condition. Simulations were generated with different numbers of replicates. The designs of two biological experiments were unbalanced with unequal replicates per condition. “No. of mod. features/site” is the number of features (i.e., peptide ions) used to estimate the abundance of a single modification. “No. of unmod. peptides/protein” is the number of peptide ions without modifications that were used to estimate the global protein abundance. “Data availability” is the ID of the MassIVE.quant repository or the GitHub repository. “Analysis” is the ID of the MassIVE.quant reanalysis container, containing analysis code and modeling results. All the experiments were conducted in data-dependent acquisition (DDA) mode.

|  | Dataset | No. of Conditions | No. of bio. replicates | No. of mod. peptides | No. of mod. features/site | No. of unmod. features/protein | Data availability | Analysis |
| --- | --- | --- | --- | --- | --- | --- | --- | --- |
| Known Ground Truth | Computer simulation 1 - Label-free | 2/3/4 | 2/3/5/10 | 1,000 | 10 | 10 |  | Github |
|  | Computer simulation 2 - Missing and low features | 2/3/4 | 2/3/5/10 | 1,000 | 2 | 10 |  | Github |
|  | SpikeIn benchmark - Ubiquitination - Label-free | 4 | 2 | 12,137 | 1.37 | 10.17 | MSV000088971 | RMSV0000000669 |
| Biological Experiment | Human - Ubiquitination - 1mix-TMT | 6 | 2 or 1 | 8,848 | 1.21 | 11.01 | MSV000088966 | RMSV0000000356 |
|  | Mouse - Phosphorylation - 2mix-TMT | 6 | 4 or 3 | 26,433 | 1.67 | 11.61 | MSV000085565 | RMSV0000000357 |
|  | Human - Ubiquitination - Label-free | 4 | 2 | 10,799 | 1.40 | 1.65 | MSV000078977 | RMSV0000000358 |

One spike-in experiment also had known changes in modified spike-in peptides, but had real-world experimental characteristics. Finally, three biological experiments demonstrated the applicability of the proposed approach across different biological organisms, modifications, experimental designs and acquisition strategies. The experimental data, R scripts with *MSstatsPTM* analysis, and results of the statistical analysis are available in MassIVE.quant (https://massive.ucsd.edu/ProteoSAFe/static/massive-quant.jsp) [8].

### Dataset 1: Computer simulation 1 - Label-free clean

#### Simulation design

The simulation represented an idealistic case. 24 synthetic label-free datasets were generated with different experimental designs and different biological variation. In each dataset, 1,000 proteins had 10 unmodified features per protein. Each of the 1,000 proteins had one PTM. Each PTM was represented by 10 modified features. The PTMs of 500 proteins had a differential fold change between conditions, while the other 500 proteins were generated with no changes in abundance between conditions. Furthermore, the fold changes of half of the 500 differential PTMs were fully masked by changes in the unmodified portion of the protein. Finally, the fold change of half the 500 non-differential PTMs was entirely due to changes in the unmodified portion of the protein. All the differential PTMs were generated with an expected log base 2 fold change of 0.75 between conditions.

Each simulation was generated with random biological variation. The observed peptide abundances were simulated by adding random noise *𝒩* (0, *σ*^2^) to the deterministic abundances described above. Two values *σ*^2^ = {.2, .3} were motivated by the experimental datasets in this manuscript.

#### Evaluation

We evaluated the ability of the statistical methods to correctly detect differentially abundant PTMs. We gauged the ability of the methods to avoid false positives (i.e. specificity), accurately estimate the fold change between conditions, and analyzed the sensitivity of detecting differentially abundant PTMs. The evaluation was performed both in the presence of confounding with changes in the unmodified protein and after applying adjustment to correct for the confounding.

### Dataset 2: Computer simulation 2 - Label-free with few low feature counts and missing values

#### Simulation design

The data were simulated as above, while providing a more realistic representation of the experiments. The feature counts and the proportion of missing values were as observed on average over all the the experimental datasets in this manuscript. Specifically, PTMs were simulated with 2 modified peptide features, and unmodified portions of the protein were simulated with 10 features. Additionally, 20% of observations for both modified and unmodified peptides were missing completely at random.

#### Evaluation

The methods were evaluated as above. We evaluated their ability to correctly detect PTM’s specificity, fold change estimation, and sensitivity. These statistics were analyzed both in the presence of, and without, confounding with the overall changes in protein abundance.

### Dataset 3: Spike-in benchmark - Ubiquitination - Label-free

#### Experimental design

**Figure 1(a)** overviews the experimental design. Four mixtures (i.e., conditions) were created with varying amounts of human lysate, background *E. Coli* lysates, and human spike-in ubpeptide mixture. Unmodified peptides from human lysate were viewed as the global proteome. Background *E. coli* lysate were used to equalize total protein levels. 50 heavy-labeled KGG motif peptides from 20 human proteins were spiked into the mixed background of the lysates. Quantitative changes in protein and site abundance of these 20 human proteins were the target of the benchmark. In particular, we distinguished the unadjusted changes (i.e. changes in the abundances of the modified peptides) and the protein-level adjusted changes of (i.e., changes in the abundances of the modified peptides relative to the changes in the abundances of the human lysate). The true log-fold changes between the relevant components of the relevant mixtures are summarized in **Figure 1(b)**. Two replicate mixtures were created per condition.

**Figure 1:**
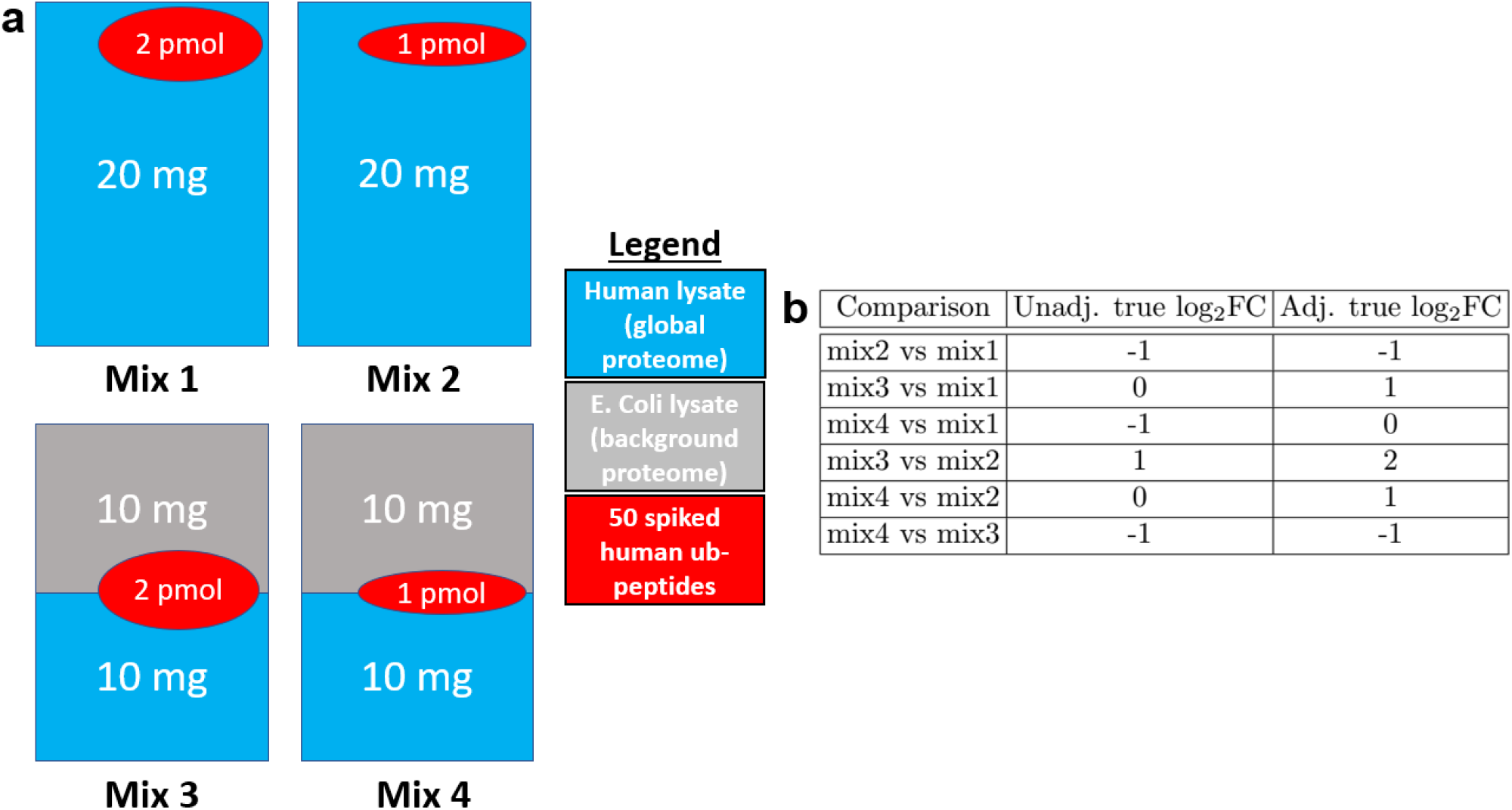
Dataset 3: Spike-in benchmark - Ubiquitination - Label-free. (a) Four mixtures (i.e., conditions) were created with varying amounts of human lysate, background *E. Coli* lysate, and human spike-in ub-peptide mixture. Unmodified peptides from human lysate were viewed as the global proteome. Background *E. coli* lysate were used to equalize total protein levels. 50 heavy-labeled KGG motif peptides from 20 human proteins were spiked into the mixed background of the lysates. Quantitative changes in protein and site abundance of these 20 human proteins were the target of the benchmark. (b) We distinguished the unadjusted changes (i.e. changes in the abundances of the modified peptides) and the protein-level adjusted changes of (i.e., changes in the abundances of the modified peptides relative to the changes in the abundances of the human lysate). “Unadj. true log_2_FC” are the log-ratios of the abundances of the spiked peptides between each condition. “Adj. true log_2_FC” was calculated by determining the ratios of the abundances of the spiked peptides and human lysate between each condition and then adjusting the ratio of the spiked peptides by the human lysate, similarly to Eq. (2).

#### Data acquisition

Each mixture was analyzed with KGG enrichment, and without KGG enrichment (i.e., in a global profiling run), with label-free LC-MS/MS. There was a 90.2% overlap of protein identifications between the identified background modified peptides and proteins quantified in the global profiling run.

#### Evaluation

We expect the relative abundances of the spike-in peptides to change as in **Figure 1(b)**. The changes in peptide abundances in all the comparisons except Mix 4 vs Mix 1 were distinct from changes in the global proteome abundances and distinct from zero, and were viewed as positive controls. In the comparison of Mix4 vs Mix 1 both the modified peptides and the global proteome background changed two-fold, and as the result the peptides in this comparison were viewed as a negative control. The background *E. Coli* lysate peptides were not expected to change in abundance in comparison, after accounting for adjustment, and were viewed as additional negative controls. We evaluated the ability of the statistical methods to avoid false positives, as well as their sensitivity in detecting the differentially abundant spike-in peptides and accurately estimate their expected fold change.

### Dataset 4: Human - Ubiquitination - 1mix-TMT

#### Experimental Design

Luchetti et al. [23] profiled human epithelial cells engineered to express IpaH7.8 under a dox inducible promoter. Uninfected cells were measured at 0 and 6 hours, while cells infected with *Shigella Flexneri* (*S. Flexneri*) bacteria were measured at 1, 2, 4, and 6 hour increments, resulting in six total conditions. 11 samples were allocated to 1 TMT mixture in an unbalanced repeated measure design. All conditions had two biological replicates except for the Dox1hr condition, which was allocated one replicate.

#### Data acquisition

The ubiquitinated peptides, and the total proteome (i.e., global profiling) were each conducted in a single LC-MS/MS run. There was a 95% overlap between the identified modified peptides and proteins that were quantified in the global profiling run.

#### Evaluation

We evaluated the ability of the statistical methods to detect changes in the abundance of modified peptides both before and after adjusting for changes in global protein abundance. The six conditions were labeled Dox1hr, Dox2hr, Dox4hr, Dox6hr, NoDox0hr, and NoDox6hr. All conditions were compared with each other, resulting in 15 pairwise comparisons. Since the dataset was a biological investigation, the true positive modifications were unknown. Shigella ubiquitin ligase IpaH7.8 was shown to function as an inhibitor of the protein Gasdermin D (GSDMD). GSDMD was actively degraded when IpaH7.8 expression was induced by dox treatment in human cells. We expect IpaH7.8 to function as an inhibitor of GSDMD in the global profiling run.

### Dataset 5: Mouse - Phosphorylation - 2mix-TMT

#### Experimental Design

Maculins et al. [24] studied primary murine macrophages infected with *S. Flexneri*. The experiment quantified the abundance of total protein and of phosphorylation in wild type (WT), and in ATG16L1-deficient (cKO) samples, uninfected and infected with *S. Flexneri*. The abundance of total protein and post-translation modifications were quantified at three time points, uninfected, early infection (45-60 minutes), and late infection (3-3.5 hours). 22 biological samples were allocated to 2 TMT mixtures in an unbalanced repeated measure design, with 11 samples allocated to each mixture. 16 replicates were spread equally between the early and late WT and cKO conditions, resulting in four replicates per condition. Both the uninfected WT and cKO contained 3 replicates, with mixture one allocating one replicate to uninfected WT and two replicates to uninfected cKO. Conversely, mixture two contained one replicate of uninfected cKO and two uninfected WT.

#### Data acquisition

This experiment included a total proteome (i.e., a global profiling run) and a phosphopeptide enrichment run. There was a 90% overlap between the identified modified peptides and proteins that were quantified in the global profiling run.

#### Evaluation

We evaluated the ability of the statistical methods to detect changes in the abundance of modified peptides both before and after adjusting for changes in global protein abundance. The six condition were labeled KO Uninfect, K_Early, KO_Late, WT_Uninfect, WT_Early, and WT_Late. 9 total comparisons were made, namely KO_Early-WT_Early, KO_Late-WT Late, KO_Uninfected-WT Uninfected, KO_Early- KO_Uninfected, KO_Late-KO_Uninfected, WT_Early-WT_Uninfected, WT_Late-WT_Uninfected, Infected- Uninfected, and KO-WT. Since the dataset was a biological investigation, the true positive modifications were unknown.

### Dataset 6: Human - Ubiquitination - Label-free no global profiling run

#### Experimental Design

Cunningham et al. [11] investigated the relationship between USP30 and protein kinase PINK1, and their association with Parkinson’s Disease. The experiment profiled ubiquitination sites, and analyzed changes in the modified site abundance. The experiment had four conditions, CCCP, USP30 over expression (USP30 OE), Combo, and Control. Cell lines were used to create two biological replicates per condition. The abundance of modified peptides was quantified with label-free LC-MS/MS.

#### Data acquisition

This experiment did not include a separate global profiling run to quantify unmodified peptides. In addition to low feature counts for unmodified peptides, this lead to substantially fewer matches between modified and unmodified peptides. There was a 41.9% overlap between the identified background modified peptides and proteins that were quantified in the global profiling run.

#### Evaluation

We evaluated the ability of the statistical methods to detect changes in the abundance of modified peptides both before and after adjusting for changes in global protein abundance. All the conditions were compared with each other in a full pairwise comparison, resulting in 6 comparisons. Since the dataset is a biological investigation, the true positive modifications were unknown.

## Background

### Goals of PTM characterization, input to statistical analyses, and notation

Consider a label-free LC-MS/MS experiment in the special case of a balanced design with *I* conditions and *J* biological replicates per condition. For simplicity, we assume that the experiment has no technical replicates, such that each biological replicate is represented by a single LC-MS/MS run. **Figure 2** schematically illustrates this data structure for one protein and one PTM site, *I* = 2 and *J* = 2. For one protein, the PTM site is represented by *K* spectral features (i.e., peptide ions, distinguished by their cleavage residues and charge states). The number of modified and unmodified features typically varies across proteins. Some log_2_-intensities can be outliers, and some spectral features can be missing. The log_2_-intensity of Feature *k*, in Replicate *j* of Condition *i* is denoted by 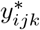. Conversely, the unmodified portion of the protein is represented by *L* spectral features, and the log_2_-intensity of Feature *l* from the unmodified portion of the protein in the same run is denoted by *y*_*ijl*_. The features can be quantified as part of a same mass spectrometry run, or in a separate enrichment and global proteome profiling run.

**Figure 2:**
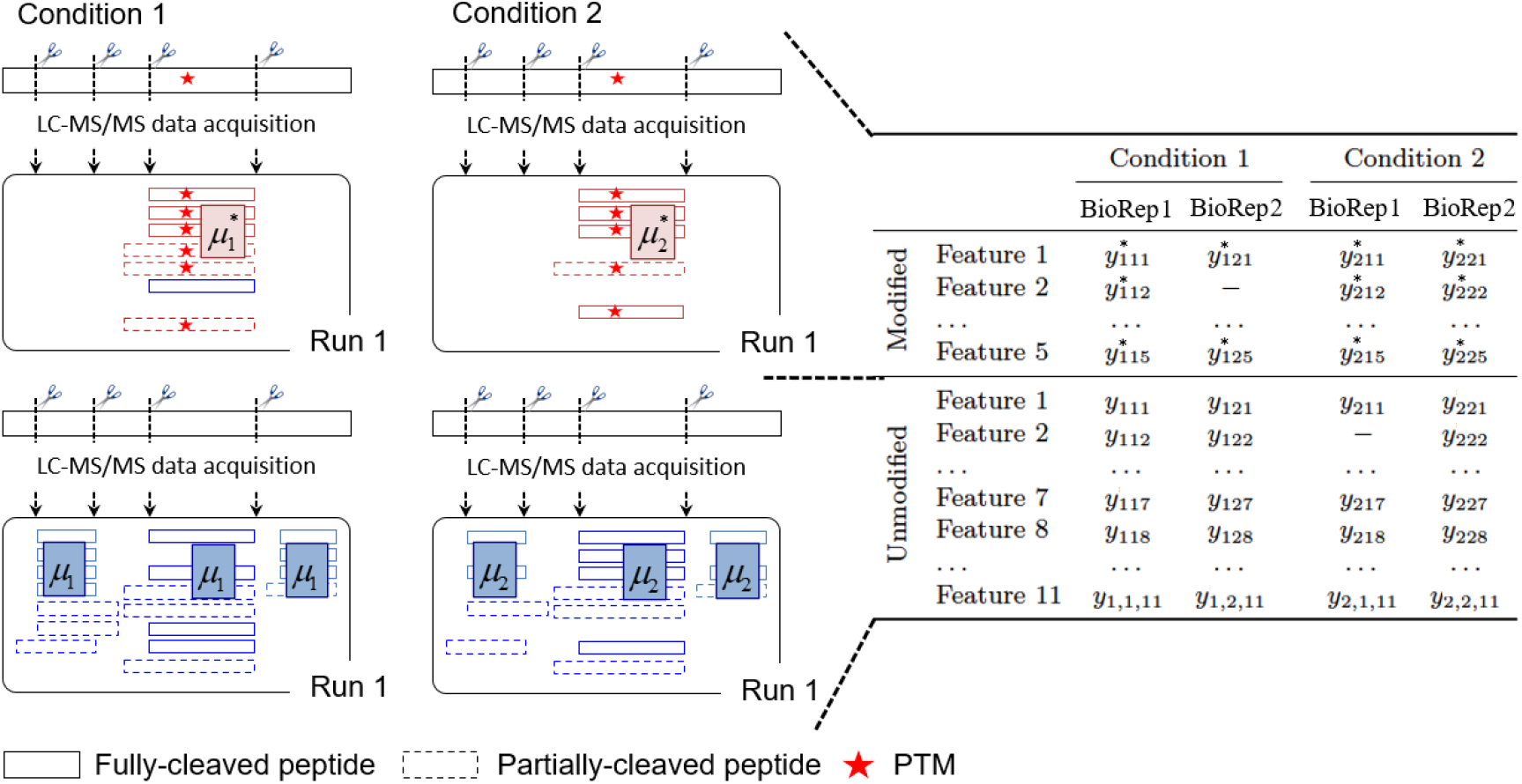
Schematic representation of one PTM site, in a special case of a label-free experiment with *I* = 2 conditions and *J* = 2 biological replicates per condition. After a log_2_ transform, we are interested in estimating the difference between the population-level PTM abundance between Condition 1 and Condition 2 (i.e, 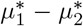), relative to the population-level difference of the overall protein abundance (i.e., *µ*_1_ *µ*_2_). These quantities are characterized by the observed spectral Features (boxes), i.e. peptides of different charge states. The peptides can be fully cleaved (solid lines), or partially cleaved (dashed lines). Unmodified features (blue) in the enriched runs are removed. The log_2_-intensities of the modified peptides in Condition *i*, Run *j*, and Feature *k* are denoted by 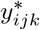. The log_2_-intensities of Feature *k* corresponding to the unmodified peptide in Condition *i* and Run *j* are denoted by *y*_*ijk*_.

The population quantity of interest is the difference between the log_2_ abundances of a PTM site in Condition *i* and Condition *i*′, denoted by 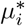 and 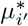 respectively. We are interested in testing the null hypothesis

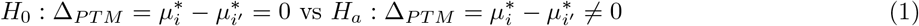

Unfortunately, this population quantity is inherently confounded with the overall changes in protein abundance. To account for this, it is advantageous to consider a different null hypothesis:

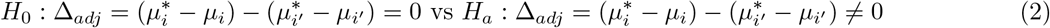

where *µ*_*i*_ and *µ*′ reflect the overall log_2_ protein abundances in Condition *i* and Condition *i*′. These quantities are estimated using protein features with and without the modification site.

### Existing statistical methods for detecting differentially abundant PTMs

#### ANOVA on summarized modified log_2_-intensities

Analysis of Variance (*ANOV A*) [21] is the simplest statistical model for summarized modified features in each biological replicate. The summarization often consists of averaging (or taking the median or other robust summary) of the log_2_ intensities of the modified features in each replicate, e.g. 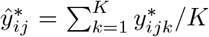. Alternatively, summarization sums the intensities of the modified features on the original scale, and then takes the log_2_

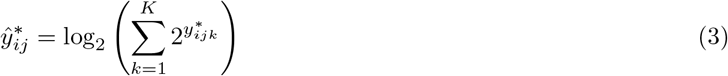

The basic *ANOV A* model is then

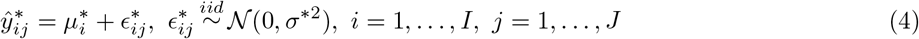

The model allows us to estimate 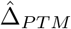 and its standard error. The estimates are used to test the null hypothesis in Eq. (1) by comparing the model-based test statistic against the Student distribution with *df* = *I*(*J* − 1) degrees of freedom in balanced designs. Unfortunately, this approach is fundamentally flawed as it does not account for the confounding between changes in the PTM abundance and the overall changes in the abundance of the unmodified portion of the protein.

#### ANOVA based on ratios of modified and unmodified log_2_-intensities

The basic *ANOV A* can be extended to account for the confounding of changes in PTM abundance and overall changes in protein abundance [34, 37, 27]. Typically this is done by first calculating sums of the intensities of the modified and unmodified features on the original scale, and then considering replicate-wise ratios of the sums and taking the log_2_

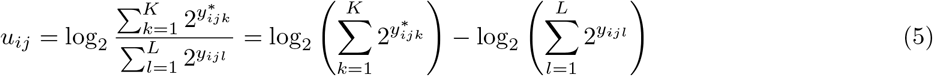

The approach then models these values with the basic *ANOV A*, which corresponds to

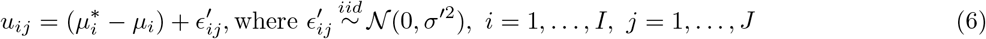

The model allows us to estimate 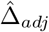 and its standard error. Based on this model, we can test the more relevant null hypothesis in Eq. (2), by comparing the test statistic against the Student distribution with *df* = *I*(*J* − 1) degrees of freedom in balanced designs.

Although effective, the approach is somewhat simplistic. It is not applicable to experimental designs with more complex sources of biological and technological variation, such as experiments with repeated measurements, experiments with multiple batches or experiments with TMT labeling. Since Eq. (5) performs the adjustment on the replicate level, the experiment must contain a matching number of replicates in both the modified and unmodified runs. Technological artifacts such as missing values further undermine the calculation of *u*_*ij*_ in Eq. (5). Finally, there is no self contained, straightforward implementation of the method, such as in the form of a coding package, and therefore the approach requires a manual implementation.

#### Limma

The estimation of nuisance variation of the two *ANOV A* models above is often further expanded with Empirical Bayes moderation implemented in *Limma* [31, 34, 35, 36, 41, 7]. *A typical application of Limma* on summarized modified log_2_-intensities takes as input 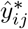, obtained as in Eq. (3), and for each PTM fits the linear model in Eq. (4). A typical application of ratio-based *Limma* takes as input *u*_*ij*_, obtained as in Eq. (5), and for each PTM fits the linear model in Eq. (6). The *Limma* versions of the models differ from the models in Eq. (4) and Eq. (6) in that they specify additional prior distributions for the model parameters. The priors are estimated from the same data by combining the information across all the proteins and all the PTM as described in [35]. Testing the null hypothesis is enhanced by combining the PTM- and protein-specific estimates of variation with a consensus estimate obtained from the estimated priors. As the result, in experiments with few biological replicates the standard errors are often smaller, and the degrees of freedom are often larger than without moderation [31]. Thus the approach tends to increase the sensitivity of detecting differential abundance.

Since *Limma* only improves upon the estimation of variation, its limitations are similar to those of *ANOV A*. In particular, the method is only directly applicable to experiments with at most two variance components, and cannot account for all the sources of variation in experiments with either isobaric labeling or complex designs. There is no self contained implementation of the methods to PTMs, requiring manual transformation and application by the user.

#### Isobar-PTM

Isobar-PTM was also proposed for experiments with LC-MS/MS quantitative strategies that employ isobaric labels such as TMT, or isobaric tag for relative and absolute quantification (iTRAQ)[5]. Isobar-PTM expresses MS measurements with a linear model and performs adjustment with respect to protein abundance using the difference between log-ratio of modified peptides in two channels and log-ratio of protein level. Unfortunately, this statistical modeling framework is not applicable to either label-free workflows or experiments with complex designs.

### Relative protein quantification in MSstats

*MSstats* [9] and *MSstatsTMT* [16] are a family of R/Bioconductor packages for statistical relative quantification of proteins and peptides in global, targeted and data-independent proteomics. The packages take as input log_2_-intensities *y*_*ijk*_. For each protein, the log_2_-intensities are first summarized into a single value per protein per run *ŷ*_*ij*_ using Tukey’s median polish [38]. The summaries are then used as input to fit a flexible family of linear mixed-effects models [26, 13, 4]. The models are fit separately for each protein. The specific model depends on the design of the experiment, labeling type and data acquisition type as summarized in **Supplementary Figure S1**. For example, the unmodified protein features in the simple design in **Figure 2** are modeled with one-way *ANOV A*

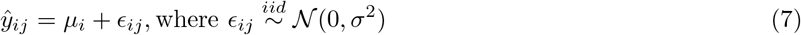

In contrast, a group comparison experiment with multiple TMT mixtures is modeled as

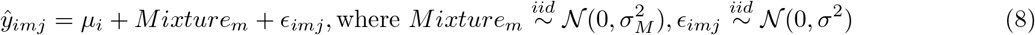

Moreover, the model fit for a particular protein depends on the pattern of missing values in that protein. If some of the terms of the model reflecting the experimental design are not estimable, a simpler model is fit for that protein instead.

Parameters of the model are estimated using restricted maximum likelihood (REML) [19]. The parameters allow us to estimate the pairwise comparison 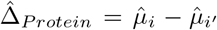 and its standard error. Similarly to *Limma, MSstatsTMT* includes an optional Empirical Bayes moderation of the standard error [16], increasing the sensitivity of detecting differential abundance when the number of biological replicates in each condition is small.

*MSstats* and *MSstatsTMT* can also be used at the feature or at the modification site level, as opposed to protein level. For example, summarizing the features per PTM site instead of per protein, the approach allows us to test the null hypothesis in Eq. (1).

The *MSstats* framework has a number of advantages over the methods above. First, unlike *ANOV A* and *Limma, MSstats* and *MSstatsTMT* are applicable to arbitrary complex experimental designs, including designs with multiple sources of variation, and unbalanced designs. Second, the approach is applicable to various data acquisition types, including label-free DDA and DIA, and experiments with TMT labeling. Third, the *MSstats* packages are compatible with various data processing tools such as Skyline, Spectronaut, MaxQuant, Progenesis, Proteome Discoverer, and OpenMS. Finally, the custom *MSstats* and *MSstatsTMT* implementation accounts for potential data artifacts, is numerically scalable and stable, and is available through both command line and a dedicated graphical user interface.

Unfortunately, the *MSstats* framework focuses on overall protein abundance, and as the result tests the null hypothesis in Eq. (1). It does not account for the confounding between the changes in PTM abundance and the overall changes in protein abundance. This manuscript proposes a simple extension to the methodology in *MSstats* and *MSstatsTMT*, to test the null hypothesis in Eq. (2).

## Results

### Statistical methods in MSstatsPTM

#### Detecting changes in PTMs, adjusted for global changes in protein abundance

The overall statistical analysis workflow and its implementation are summarized in **Figure 3**. *MSstatsPTM* takes as input the modified spectral features 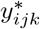, and the corresponding unmodified features *y*_*ijk*_. Ideally, the modified features are acquired separately after an enrichment, to maximize the information content in the resulting dataset, and the unmodified features are acquired separately as part of a global proteome profiling. However the method can also take as input a combination of modified and unmodified features acquired within a same run.

**Figure 3:**
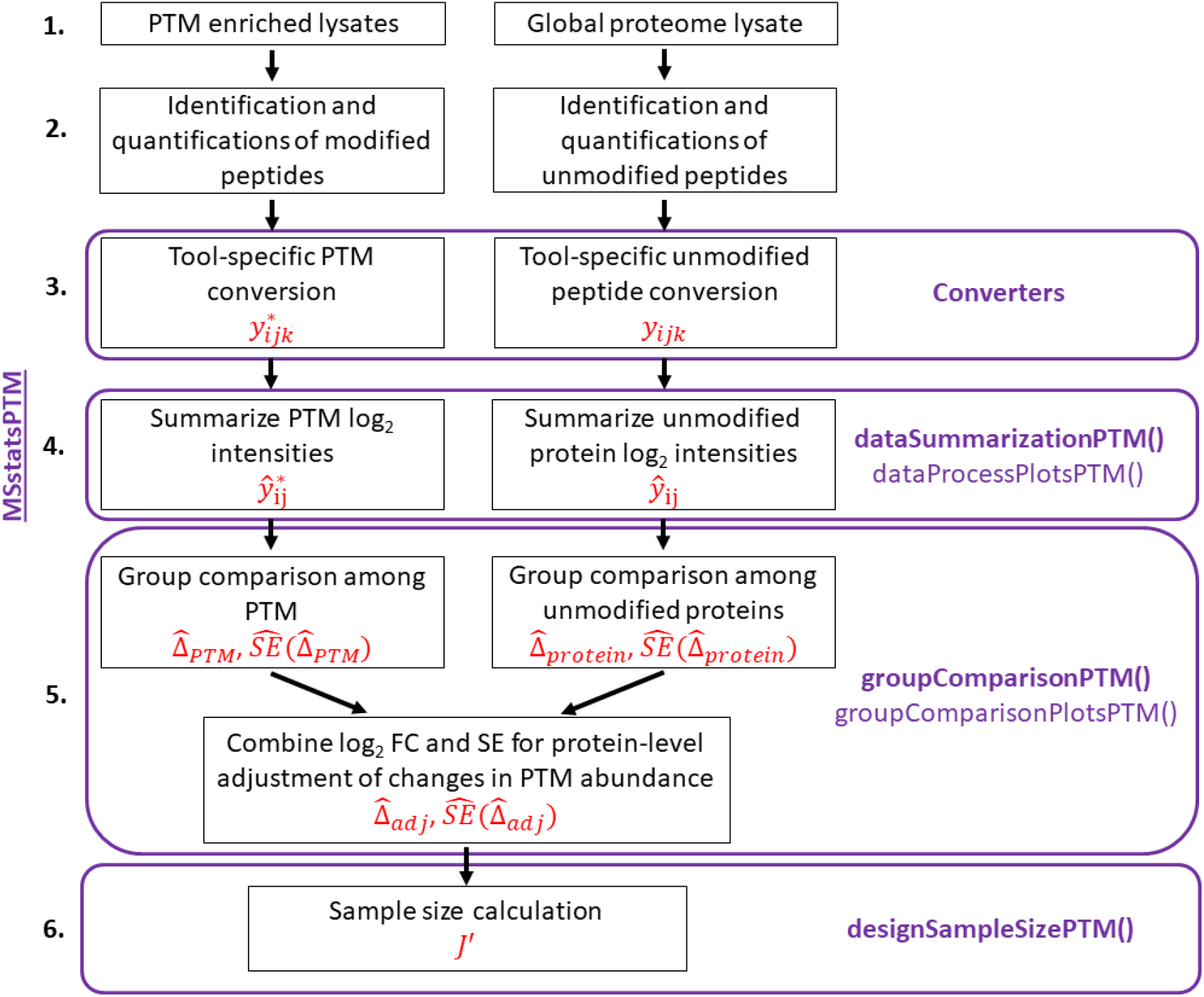
The *MSstatsPTM* workflow. The names of the *MSstatsPTM* R functions used for each step are highlighted in purple and the output notations are highlighted in red. The workflow begins with the acquisition of the enriched and global proteome lysates. The package is applicable to label-free data acquisitions such as DDA, DIA, SRM and label-based data acquisitions such as TMT. It takes as input lists of identified and quantified spectral features for the PTM and for the unmodified portion of the protein, produced by spectral processing tools such as MaxQuant, Progenesis, or Spectronaut. Conversion, summarization and statistical modeling are performed separately for the PTM and for the unmodified portions of the proteins. Steps 4 and 5 leverage the summarization and modeling functions from *MSstats* and *MSstatsTMT*. Modelbased summaries are combined to adjust the changes in the PTM abundance for changes in abundance of the unmodified portion of the protein. Finally, sample size calculation for future experiments can be performed using the modeling output.

Each feature type is first analyzed separately, with *MSstatsPTM* methods calling the relevant functionalities in *MSstats* (for label-free experiments) or *MSstatsTMT* (for experiments with TMT labels). In particular, the modified features are summarized into run-level summaries 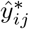.

The estimated summaries of the modified features are used as the input to models such as in Eq. (7) or Eq. (8). The resulting model-based estimates include 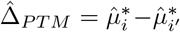, and its standard error 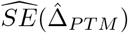. Similarly, the unmodified features of each protein are summarized for each run into ŷ_*ij*_, and the summaries are used as input to a separate analysis by *MSstats* or *MSstatsTMT* producing 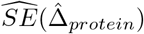. From these summaries, the proposed approach estimates the adjusted difference 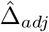 in Eq. (2)

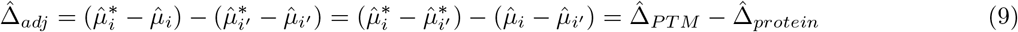

Assuming that the sources of variation in the summaries of modified features that are unexplained by the model are independent from the sources of variation in the summaries of modified features, the standard error 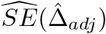 is obtained by combining the standard errors from the two model fits

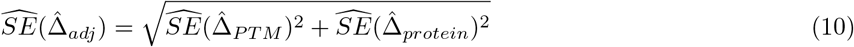

For example, in the simple case of **Figure 2** with *J* = 2 replicates, where 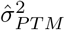 and 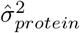 are respectively the estimates of the error variance for the PTM and protein model described in Eq. (7), the standard error is calculated as

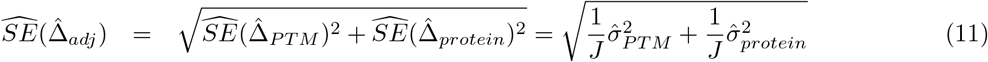

The estimated standard error is larger than the standard errors associated with each individual feature type, reflecting the combined uncertainty in the two estimates. Finally, the degrees of freedom associated with Eq. (10) are obtained via the Satterthwaite approximation [21, 33]

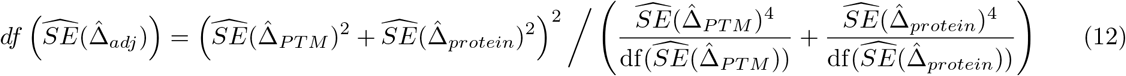

To test the null hypothesis in Eq. (2), the test statistic 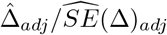 is compared with the Student distribution with the degrees of freedom in Eq. (12). The p-values of the comparison are adjusted for multiple testing using the approach by Benjamin and Hochberg [2].

#### Sample size calculation for future PTM experiments

The statistical framework in *MSstatsPTM* enables sample size calculation for future experiments studying changes in PTM. The procedure has been described in general in [21], and for protein significance analysis specifically in [28]. It requires us to specify the desired levels of the following quantities: a) *q*, the False Discovery Rate of detecting differential abundance, b) *β*, the average Type II error rate, c) ∆_*adj*_, the minimal log_2_-fold change in adjusted PTM abundance of interest, d) *m*_0_*/*(*m*_0_ +*m*_1_), the fraction of truly differentially modified PTM sites in the comparison, and e) 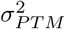 and 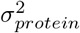, the anticipated variances associated with the pairwise comparisons of conditions for the modified and unmodified protein summaries, respectively. Typically, these variances are estimated from an existing experiment, conducted with the same biological material and measurement workflow.

Given the above quantities, and assuming a balanced design that allocates the same number of replicates *J* ′ to both modified and unmodified profiling runs, the minimal number of replicates *J* ′ for each of *I* conditions is chosen to bound the variance of the estimated log_2_-fold change SE^2^(∆_*adj*_):

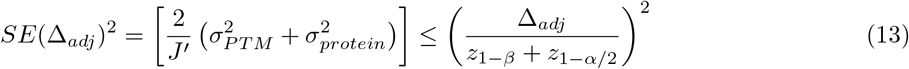

Where

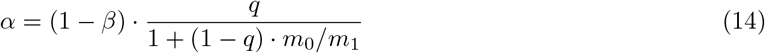

and *z*_1*−β*_ and *z*_1*−α/*2_ are the 100(1 − *β*)^th^ and the 100(1 − *α/*2)^th^ percentiles of the Standard Normal distribution. Solving for *J* ^*i*^, the number of biological replicates per condition is

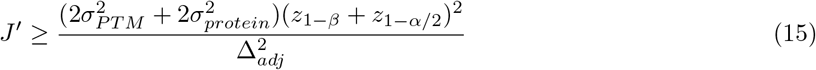

The numerator has two sources of variation, reflecting a larger uncertainty in the adjusted calculation. Therefore, the adjustment typically requires a larger sample size to gain the same sensitivity as the unadjusted estimation.

In Eq. (15) we calculated the required sample size, however in some cases the statistical power of an experiment may be of interest. In this case, Eq. (13) is still used, however the number of replicates, *J* ′, is fixed and False Discovery Rate, *q*, is solved for.

### Implementation of MSstatsPTM

The implementation of the open source R package *MSstatsPTM* is overviewed in **Figure 3**. By leveraging the implementations in *MSstats* and *MSstatsTMT*, the proposed approach is versatile. It is applicable to a wide variety of experimental designs, including group comparison, paired designs, time course designs and unbalanced designs. It is applicable to label-free data acquisitions such as DDA, DIA, SRM and label-based data acquisitions such as TMT. It can model experiments where the experimental designs for PTM profiling and global proteome profiling vary in properties such as number of biological replicates, data acquisition strategies and runs.

*MSstatsPTM* takes as input lists of identified and quantified spectral features, produced by spectral processing tools such as MaxQuant, Progenesis, or Spectronaut (Step 3 of **Figure 3**). Conversion is performed separately for the runs enriched in modified peptides, and separately for the global profiling runs. We require the processing tools to identify the modification site (i.e., the amino acid in the protein sequence where the modification occurred). This will generally include the amino acid abbreviation, plus its number in the protein sequence. For example, a modification on a 70th amino acid in the sequence, serine should be marked as “S70”. Occasionally the outputs of data processing tools only include the peptide sequence with the modified amino acid highlighted, without indicating the location in the protein sequence. For these cases *MSstatsPTM* includes functionality for identifying the location, given the modified peptide sequence and a FASTA file with the entire protein sequence. The converters output the modified spectral features 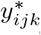, and the corresponding unmodified features *y*_*ijk*_ in the format required for summarization.

The next step is PTM/protein summarization using the *dataSummarizationPTM* () function (Step 4 of **Figure 3**). Summarization is performed separately for the PTM features, and for the features representing the unmodified portion of the protein. When summarizing the PTM, modified peptide features that span the same modification site are summarized together. Peptides that include multiple modifications are not included in the single modification summarization, and are grouped separately. The unmodified protein summarization is performed as discussed above for *MSstats*. When summarizing the unmodified protein features, the package optionally imputes missing values using an Accelerated Failure Time (AFT) model [39]. When summarizing the modified features, missing value imputation is also possible but should be performed with care. PTMs generally exhibit low feature counts and may be missing due to reasons other than low abundance. These issues can violate the assumptions underlying the imputation, and lead to numerically unstable results. The outputs of this step are the run-level summaries for the modified 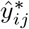, and unmodified ŷ_*ij*_, features.

Separate statistical models are fit to both feature summaries using the *groupComparisonPTM* () function (Step 5 of **Figure 3**). *MSstats* or *MSstatsTMT* models are leveraged to automatically reflect the experimental design and the data acquisition. If the base model is not applicable for a particular PTM or protein, e.g. due to missing data, a simplified model is fit. The output of the models are the estimates 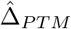 and 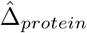, and their standard errors 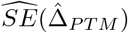 and 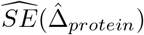.

After modeling, the model for the modified peptides is adjusted for changes in the abunance of the unmodified portion of the protein, using the methods described above. Modification sites which lack corresponding global profiling information cannot be adjusted for changes in protein abundance. In this case the implementation reverts to testing the null hypothesis in Eq. (1) using the statistical methods seen in *MSstats*, applied separately to each modified peptide. The final output is the estimate 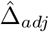 and it’s standard error 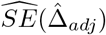.

Finally, the statistical models can be used to calculate the required sample size for future PTM experiments using the *designSampleSizePTM* function (Step 6 of **Figure 3**).

In addition to the above functionalities, the implementation includes visualizations for quality control, *dataProcessPlotsPTM* (), and assessment of the quality of model fit, *groupComparisonPlotsPTM* ().

The implementation relies on functionalities from the R packages *MSstats* [9] and *MSstatsTMT* [16], which in turn rely on the R packages *lme*4 [1] and *lmerTest* [22]. *MSstatsPTM* is available on Bioconductor, http://www.bioconductor.org/packages/release/bioc/html/MSstatsPTM.html, and Github, https://github.com/Vitek-Lab/MSstatsPTM.

### Evaluation

#### Evaluation criteria

We compared the performance of *MSstatsPTM* to that of *Limma* and *ANOV A*, both before and after adjusting for changes in the unmodified portion of the protein. Since *Isobar*-*PTM* is only applicable to experiments with TMT labeling it could not be applied to the datasets with known ground truth in this manuscript, and was therefore excluded from the comparisons.

*MSstatsPTM* before adjusting for changes in the unmodified portion of the protein corresponds to base *MSstats* or *MSstatsTMT*, modeled on the peptide level as described in **Supplementary Figure S1**, as appropriate for the experimental design. *MSstatsPTM* with the adjustment described in **Section 3** was used without imputing missing values, and without Empirical Bayes moderation.

*Unadjusted ANOV A*, i.e. *ANOV A* before adjusting for changes in the unmodified portion of the protein, was as in Eq. (4). *Adjusted ANOV A*, i.e. *ANOV A* with the adjustment was modeled as in Eq. (6). Finally, *unadjusted Limma* used the same model formula in Eq. (4), while including a moderated variance estimation. *Adjusted Limma* was modeled as in Eq. (6) including moderated variance estimation. All the evaluations were done at the FDR-adjusted p-value cutoff of *q* = .05. More details are in **Supplementary Sec. 3**.

We evaluated *MSstatsPTM* on simulated and spike-in datasets with known ground truth in terms of true positives (*TP*), false positives (*FP*), true negatives (*TN*), and false negatives (*FN*) differentially abundant PTMs. The true positives were defined as PTMs with changes distinct from the overall changes in abundance of the unmodified portion of the protein. The true negatives were defined as PTMs which, after accounting for the changes in the overall protein abundance, were not differentially abundant. Additional summaries were performed including accuracy, recall, and positive predictive value (PPV)/empirical False Discovery Rate (eFDR) as described in Eq. (16).

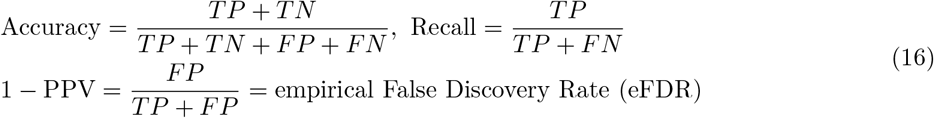

For biological experiments with unknown ground truth, we compared the differentially abundant PTMs with and without adjusting for changes in unmodified protein abundance.

#### Protein-level adjustment was required to control eFDR in differentially abundant PTM

**Figure 4(a)** summarizes the eFDR reported on Computer Simulation 1 dataset by *MSstatsPTM, adjusted ANOV A, adjusted Limma* and base *MSstats, unadjusted ANOV A* and *unadjusted Limma* methods. The simulation mimicked a “clean” label-free group comparison experiment, not compromised by issues such as deviations from model assumptions, missing values and outliers. All the analyses were performed to control the eFDR at at most 5%. Yet, even under these favorable circumstances, the models that did not adjust for confounding from changes in overall protein abundance produced an excessive number of false positives. The versions of the models that accounted for the confounding produced error rates that were much better calibrated at the desired level.

**Figure 4:**
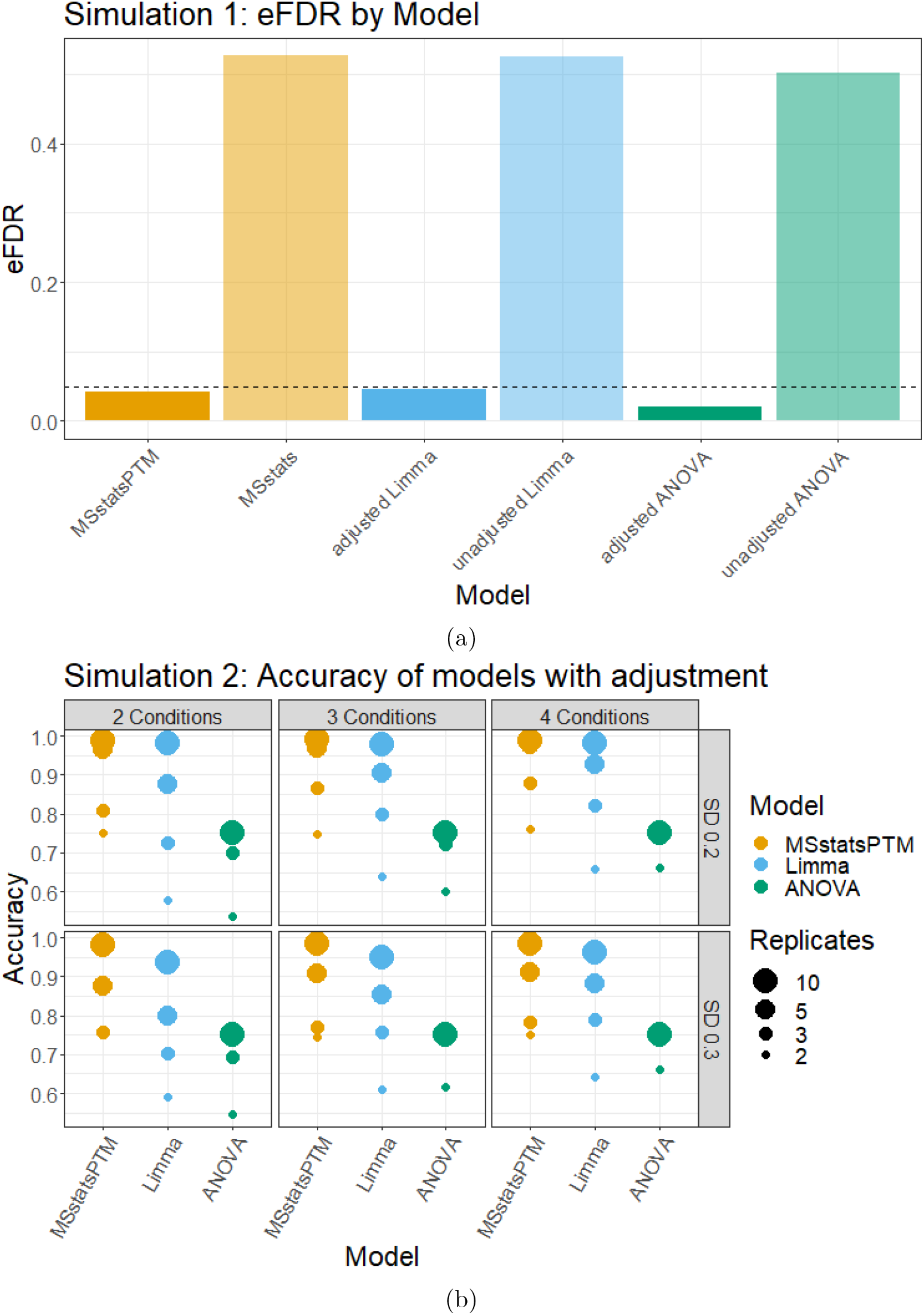
Computer simulations. a) Dataset 1, clean simulation, analyzed to control eFDR at at most 5% (horizontal line). The methods not accounting for the confounding between changes in PTM and overall changes in protein abundance produced exceedingly high numbers of false positives. In contrast, the methods accounting for the confounding correctly calibrated the proportion of false positive differentially abundant PTM. b) Dataset 2, a noisy simulation, which included limited feature observations and missing values. *MSstatsPTM* had higher accuracy than *ANOV A* and *Limma* given the same noise, number of biological replicates, and number of conditions.

#### In noisy simulations, MSstatsPTM more accurately detected differentially abundant PTM

**Figure 4(b)** summarizes the overall accuracy of methods adjusting for changes in abundance of the unmodified portion of the proteins in Computer Simulation 2. The simulation mimicked a more realistic label-free group comparison experiment, including low counts of modified features and missing values. We evaluated the impact of low versus high noise, of the number of biological replicates, and of the number of conditions. For low noise, *MSstatsPTM* outperformed the existing methods across all conditions and number of replicates, with near 100% accuracy when the replicates were high. As the noise increased the accuracy of all the methods decreased, however *MStatsPTM* still outperformed the existing methods. The difference was primarily due to two reasons. First, the ratio-based summarization approach used by *Limma* and *ANOV A* requires measurements for both the PTM and unmodified protein. In contrast, *MSstatsPTM* can leverage the information in the PTM or unmodified portion of the protein if one of the two is missing. On average over all simulations, 3.94% of the total unmodified protein run summarizations used by *MSstatsPTM* were discarded by *Limma* and *ANOV A* due to missing PTM data. No PTM run summarizations were discarded due to missing protein data. Second, the robust Turkey Median Polish summarization in *MSstatsPTM* was more resistant to outliers.

**Supplementary Figure S2** further compares the fold change estimation across all modified peptides. *MSstatsPTM* showed a tighter distribution of estimated fold changes around the true fold change. Specifically, the inter-quartile range (IQR) of the estimated fold change for *MSstatsPTM* was on average 32.5% smaller than *Limma* and *ANOV A*’s IQR. While the mean of the estimated fold changes was generally correct for all the methods, the proposed approach correctly estimated the fold change more often across all PTMs.

#### In the label-free benchmark experiment, MSstatsPTM had a higher sensitivity

**Figure 5(a)** summarizes the evaluation on the label-free spike-in group comparison experiment, for all the methods, with and without adjusting for changes in abundance of the unmodified portion of the protein. Without adjusting for changes in unmodified protein abundance, all the approaches incorrectly estimated the log_2_-fold change of the modified spike-in peptides. After adjustment, the estimation was generally in line with the ground truth for all methods, however *MSstatsPTM* ‘s distribution of estimated fold changes was tighter. On average over all comparisons the inter-quartile range of the estimates by *MSstatsPTM* was 32.86% smaller than that by *Limma* and *ANOV A*. As seen in the previous section, the robust summarization by *MSstatsPTM* was particularly useful when the number of features was low. This was also the case in this experiment, with an average of 1.37 features per PTM. Additionally, the ratio based summarization used by *Limma* and *ANOV A* discarded more data than in the simulations. The ratio based summarization lost 9.63% of the unmodified protein and 1.94% of the PTM run summarizations used by *MSstatsPTM*. This additional information lead *MSstatsPTM* to a more accurate fold change estimation and better calibrated variance.

**Figure 5:**
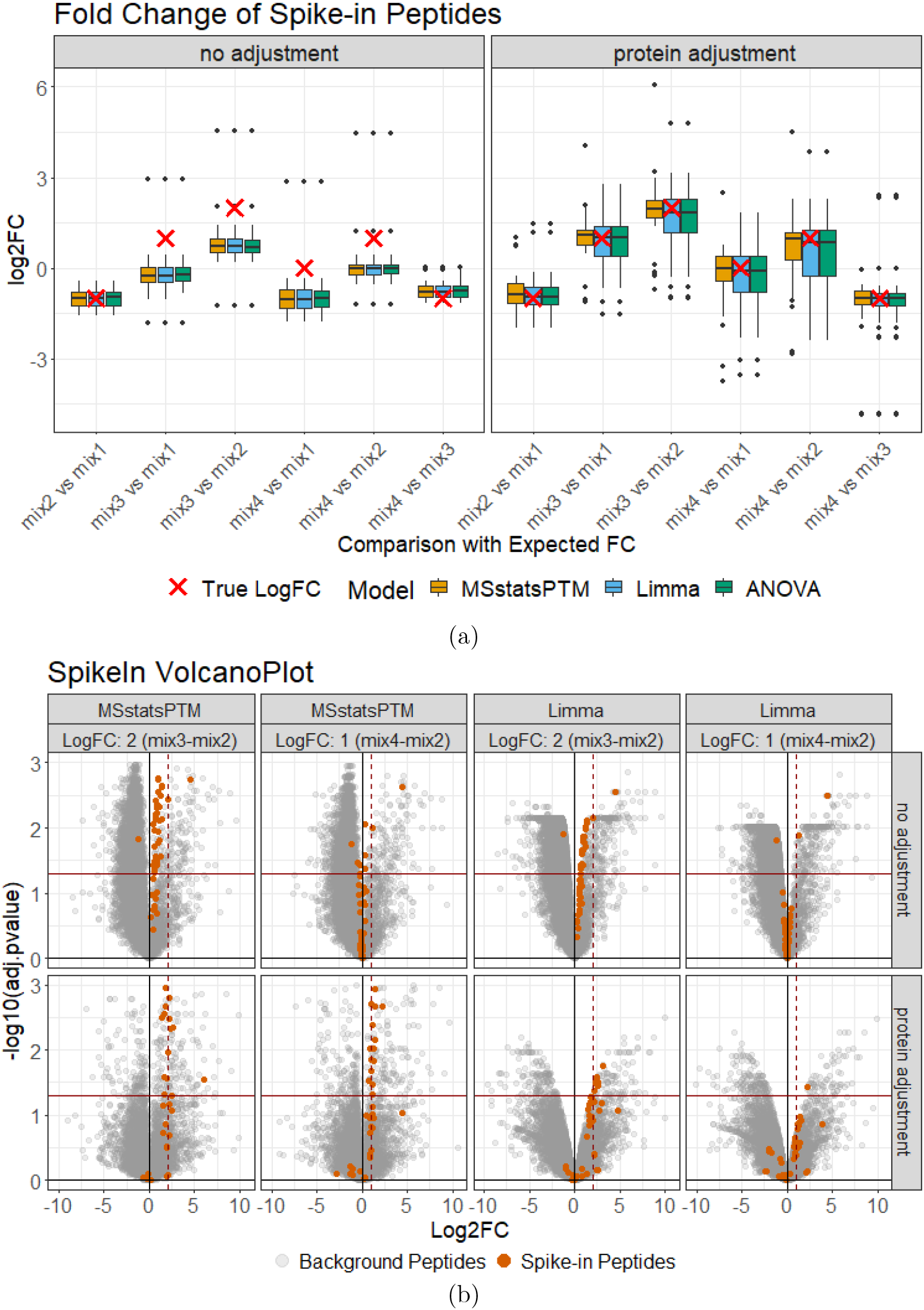
Dataset 3: Spike-in benchmark - Ubiquitination - Label-free. a) The distribution of spike-in peptides log_2_ fold change estimated by *MSstatsPTM, Limma*, and *ANOV A* with and without adjustment. The expected log_2_ fold change is highlighted by a red ‘X’. Protein adjustment removed systematic differences from the expected log_2_ fold change in all models. b) The statistical results of *MSstatsPTM* and *Limma* modeling the *mix*3-*mix*2 and *mix*4-*mix*2 comparisons before and after adjustment. The solid horizontal line shows the adjusted pvalue cutoff of .05. The solid vertical line shows log_2_ fold change of 0. The dashed vertical line shows the expected log_2_ fold change of the spike-in peptides. Before adjustment the spike-in peptides did not follow the expected log_2_ fold change, but were more in line with expectation after adjustment.

**Figure 5(b)** details the detection of differentially abundant PTM for *MSstatsPTM* and *Limma*, with and without the adjustment for changes in abundance in the unmodified portion of the protein, for the *mix*3-*mix*2 and *mix*4-*mix*2 comparisons. As above, the log_2_-fold changes of the spike-in peptides were only correctly estimated when accounting for changes in the unmodified protein abundance. Additionally, the background peptides, serving as the null model, show many false positives before adjustment. After adjustment the extent of false positives substantially decreased. Specifically, for *MSstatsPTM* the number of false positives went from 20.88% to 1.84% after the adjustment, and for *Limma* went from 26.04% to 1.18%. While the proposed method and *Limma* both correctly estimated the fold change of the spike-in peptides, using *Limma* resulted in many large adjusted p-values, and lower sensitivity. This was mainly due to *Limma* estimating a higher variance, even when including variance moderation. On average, over all the PTMs in this experiment, the variance components estimated by *Limma* were 35.7% larger than for *MSstatsPTM*. Volcano plots for all methods and comparisons can be seen in **Supplementary Section 3.2**.

#### In two biological experiments with TMT labeling, MSstatsPTM corrected for confounding between changes in the PTM and changes in the unmodified protein

**Figure 6(a)** summarizes the results of Dataset 4: Human - Ubiquitination - 1mix-TMT in terms of number of differentially abundant PTMs before and after adjustment. Adjusting for changes in abundance in the unmodified portion of the protein caused fewer PTMs to be detected as differentially abundant. A question is whether this was due to adjustment in the log_2_-fold estimation, or to an increase in standard errors during the adjustment (Eq. (10)). To check that, we considered modified peptides for which the adjusted log_2_-fold change was within 10% of the unadjusted log_2_-fold fold change, but which lost statistical significance after the adjustment. In the case of Dataset 4, only one PTM became non-differentially abundant due to an increase in standard error. In other words, the decrease in differentially abundant PTMs was primarily due to removing the confounding with global protein abundance, and not to larger variance estimates.

**Figure 6:**
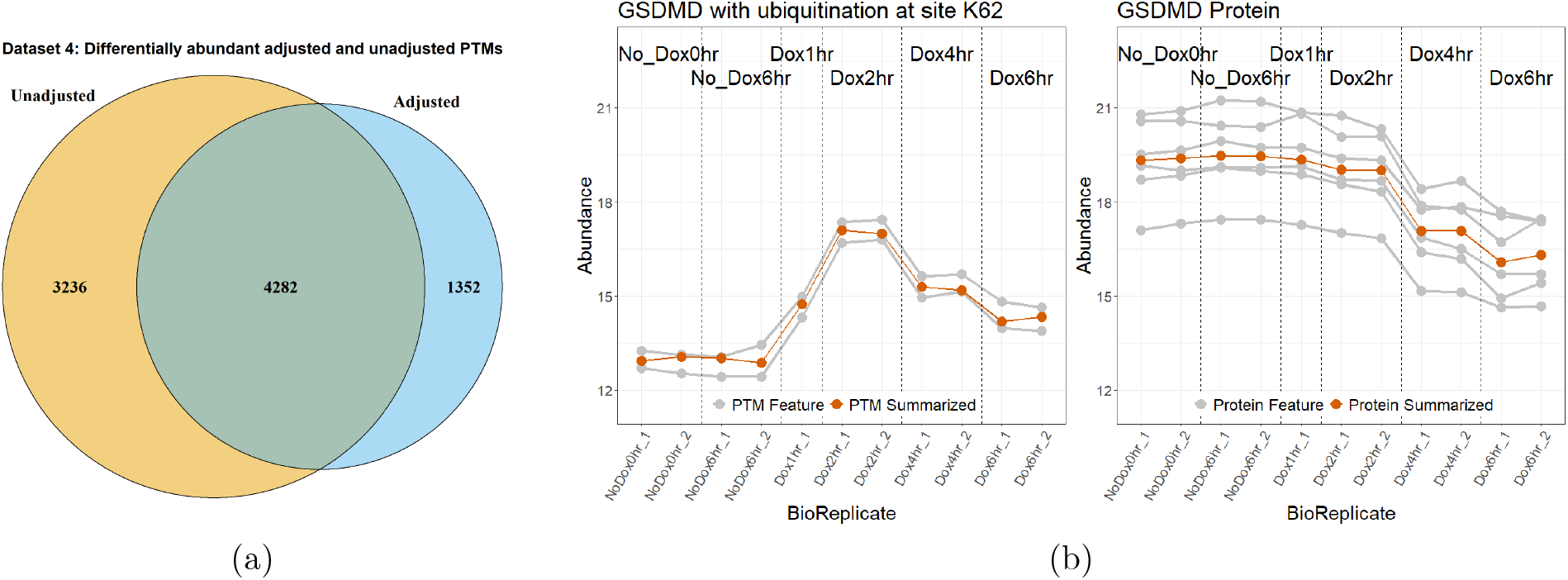
Dataset 4: Human - Ubiquitination - 1mix-TMT, analysis with *MSstatsPTM*. (a) The overlap of differential modified peptides for the PTM model with and without global protein level adjustment across all pairwise comparisons. (b) Comparing the profiling of protein *GSDMD* with the ubiquitination at site *K*62. The individual PTM and protein features are shown in grey, while the summarization is highlighted in red.

Cases where true changes in PTMs were masked by changes in abundance in the unmodified portion of the protein were less frequent. One such case is in **Figure 6(b)**. Luchetti et al. [23] showed that *GSDMD* was actively degraded when IpaH7.8 expression was induced by Dox treatment. Our reanalysis confirmed that the *GSDMD* protein was down-regulated when Dox treatments reached the 4 and 6 hour marks. Conversely, ubiquitination of *GSDMD* at site *K*62 up-regulated abundance between the same conditions. This up-regulation was originally confounded by the down-regulation of unmodified *GSDMD*, and made the modification appear to have little change between no Dox and Dox 4 and 6 hour conditions. The proposed approach accounted for this confounding and the modification was detected as differentially abundant, with a log_2_ fold change of 2.79 between the Dox 1 hour and Dox 4 hour conditions (**Supplementary Figure S6**). The change in PTM abundance would have been challenging to observe without the proposed approach. Indeed, the modification contradicts the previous research focusing on global profiling of the *GSDMD* protein [23].

**Figure 7(a)** illustrates a similar result for Dataset 5: Mouse - Phosphorylation - 2mix-TMT. Adjusting for changes in abundance in the unmodified portion of the protein caused fewer PTMs to be detected as differentially abundant. As above, we checked whether this change was due to fold change adjustment or an increased standard error. In the case of Dataset 5, 548 PTMs were not detected as differential abundant due to an increased standard error. This corresponded to 3.4% of all the PTMs that became non-differentially abundant.

**Figure 7:**
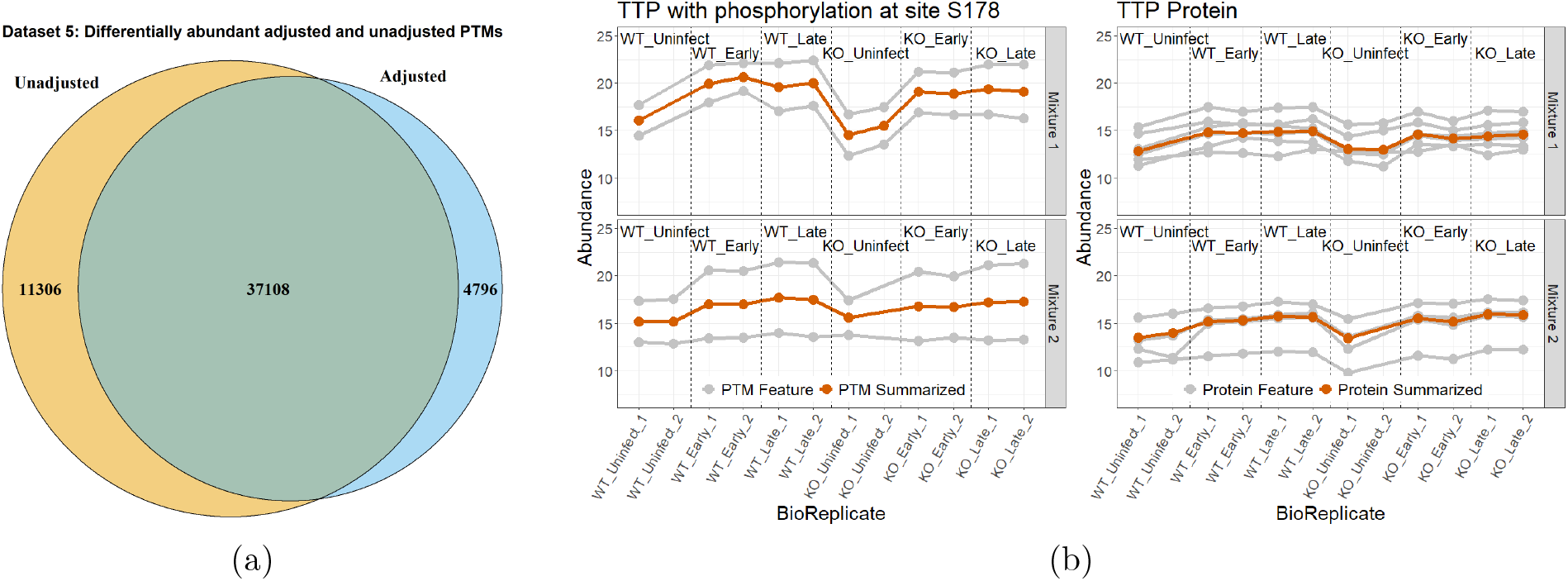
Dataset 5: Mouse - Phosphorylation - 2mix-TMT, analysis with *MSstatsPTM*. (a) The overlap of differential modified peptides for the PTM model with and without global protein level adjustment across all pairwise comparisons. (b) Comparing the profiling of protein *TTP* with the phosphorylation at site *S*178. The individual PTM and protein features are shown in grey, while the summarization is highlighted in red. The plots are separated according to TMT Mixtures 1 and 2.

**Figure 7(b)** shows an opposite case where the protein exhibits a change in abundance while the modification does not. Before adjusting for changes in the unmodified protein, the modification at *S*178 of protein *TTP* was shown to be differentially abundant between WT Uninfect and WT Late, with a log_2_ fold change of 2.9. However the unmodified protein was shown to contribute 69.5% of this change, while the modification only accounted for 30.5% after adjustment (**Supplementary Figure S7**). This caused the modification to lose differential abundance.

#### In label-free experiment without a separate global profiling run, MSstatsPTM eliminated the confounding due to changes in the unmodified protein, albeit less effectively than in the presence of a global profiling run

Unlike the other datasets in this manuscript, Dataset 6: Human - Ubiquitination - Label-free had no unmodified global profiling run. Therefore, after peptide identification and quantification, data from unmodified peptides were used separately in place of a global profiling run. This resulted in a sparse coverage of the modified features by the unmodified protein counterparts. Of the 10,799 identified ubiquitination sites, only 4,526 had features from unmodified portion of the same protein. A PTM without features from the unmodified portion of the protein could not be adjusted for the confounding. Additionally, the lack of a separate global profiling run resulted in low feature counts and noisier measurements of the unmodified peptides as compared to the other experiments (Table 1).

Figure 8. shows the number of differentially abundant PTMs before and after adjustment. After adjusting for changes in abundance in the unmodified portion of the protein, the number of differentially abundant PTMs decreased. However this was mainly due to the lack of a global profiling run. The decrease in differentially modified PTMs was smaller among PTMs with available unmodified protein counterparts. As above, we tested if this drop in differentially abundant PTMs was due to an increase in standard error. Here only 25 PTMs lost differential abundance due to increased standard error.

**Figure 8:**
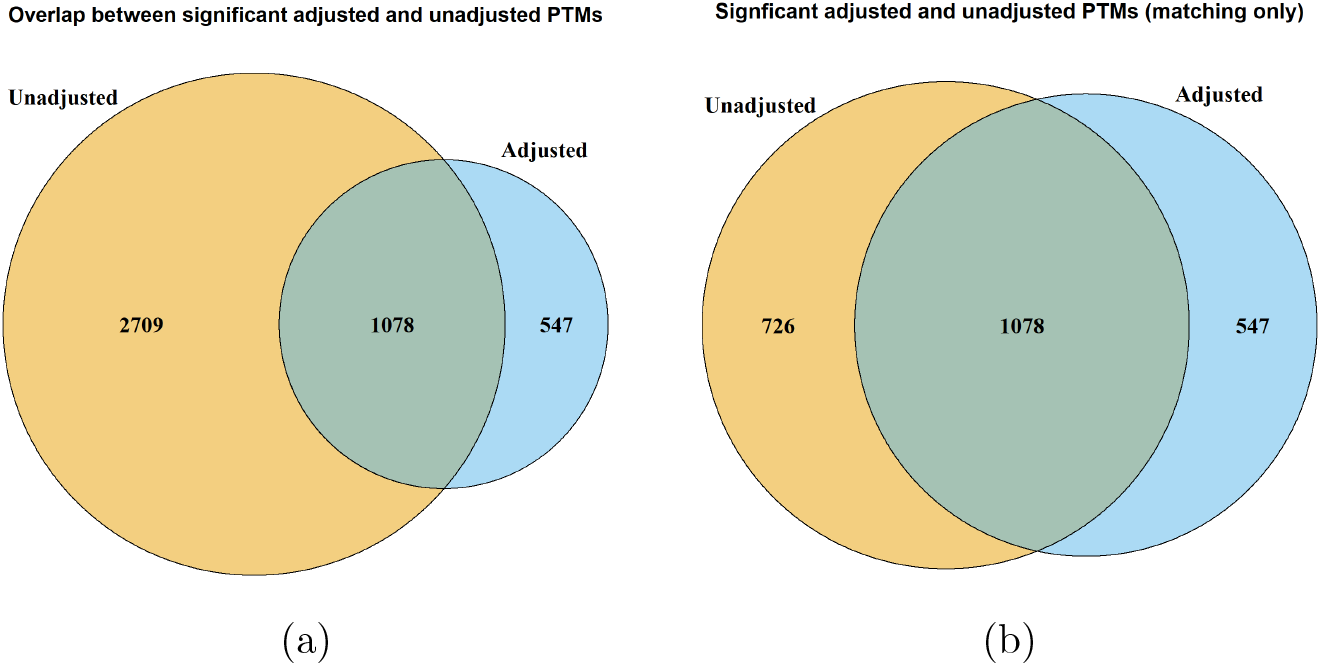
Dataset 6: Human - Ubiquitination - Label-free no global profiling run, analysis with *MSstatsPTM*. a) The overlap of differential modified peptides for the PTM model with and without global protein level adjustment across all comparisons, including all measured PTMs. b) The overlap of differential modified peptides with and without global protein level adjustment, including only PTMs for which features from the unmodified portion of the same protein were also available.

#### Noisy PTM measurements benefited from additional biological replicates

Sample size calculations and statistical power analyses help evaluate the benefit of additional biological replicates, in particular when measurements are noisy. Since the proposed approach does not have the restriction of the equal number of modified and unmodified global profiling runs, it is also interesting to evaluate the interplay between the number of replicates of each type in presence of differing amounts of noise. In Datasets 4 and 5 the variance of the PTMs was higher than that of the unmodified protein summaries, with median values of PTM variance of .45, and protein variance of .3. In Dataset 6 the variance of the PTM and the unmodified protein summaries were comparable, with a median variance of .45. The power analysis took as input these median variance values.

Figure 9. shows that regardless of the relative amount of variation, larger adjusted log_2_-fold changes and larger number of replicates enabled larger statistical power. When the variance of PTMs was equal to the variance of global protein profiles, it did not matter whether we increased the number of PTM runs or the number of unmodified protein runs. However, when the variance of the PTM summaries was higher than that of the unmodified protein summaries, allocating more biological replicates to the PTM profiles lead to a more efficient increase of statistical power.

**Figure 9:**
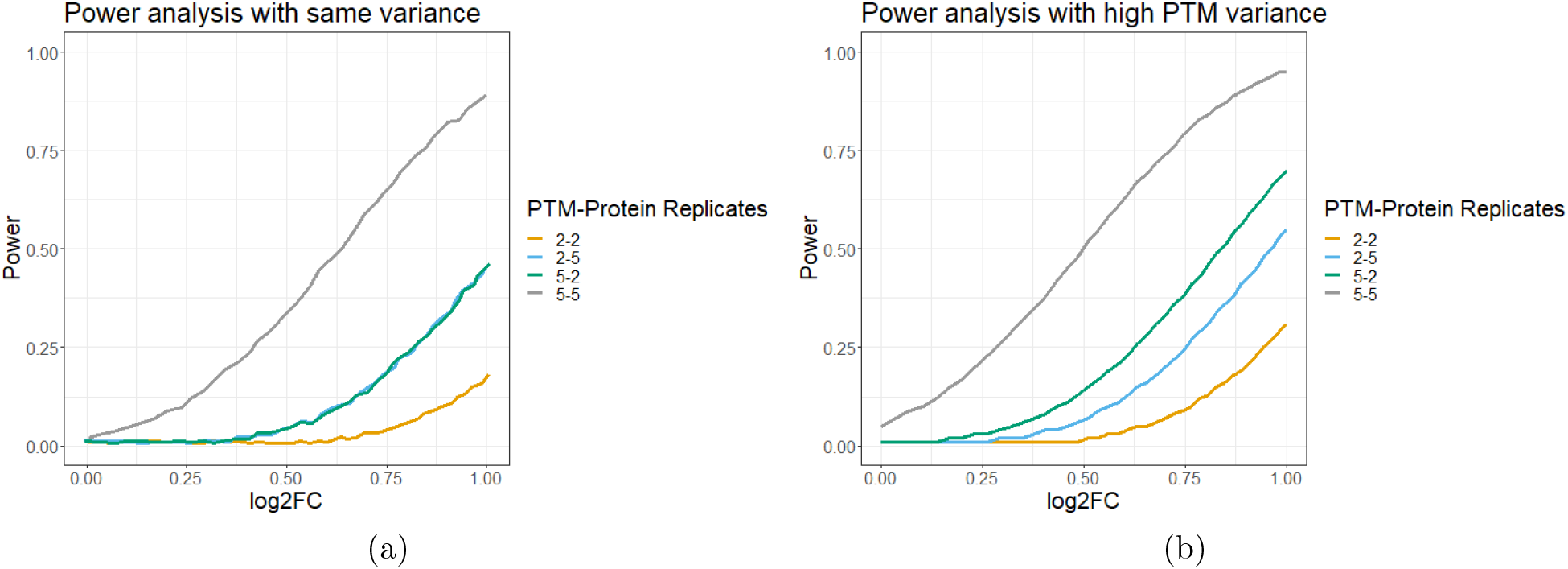
Power analysis with with *MSstatsPTM*, based on variance components from Datasets 4, 5, & 6. Y-axis: statistical power, i.e. the probability of rejecting the null hypotheses of no protein-adjusted change in a PTM between conditions, when in fact the change exists. X-axis: log_2_ fold change, after for adjusting for changes in the unmodified protein. a) The power of detecting differential abundant PTMs, when the modified features and the unmodified protein summaries have the same variance .45. b) The power of detecting differential abundant PTMs, when the modified features had a larger variance (.45) as compared to the unmodified protein summaries (.3).

## Discussion

We proposed a statistical modeling framework for detecting differentially abundant PTM, and its implementation in *MSstatsPTM*. The proposed approach removes the confounding of changes in PTMs with changes in the unmodified protein summaries, and can reveal modifications of interest that are otherwise entirely masked by changes in abundance in the unmodified portion of the protein. This is valuable, because many PTMs are often found non-differentially abundant prior to the adjustment, and validating them manually to establish false negatives is generally unfeasible.

Our results show that *MSstatsPTM* is more accurate and more sensitive than the existing approaches. The gain is due to a more efficient use of the data, and to a more accurate representation of the systematic and random variations. While the ratio-based approaches of *ANOV A* and *Limma* first consider differences within samples, *MSstatsPTM* first considers the differences between conditions. This enables a greater flexibility in terms of modeling complex designs, accounting for outliers and missing values, and planning subsequent experiments.

The implementation of *MSstatsPTM* is a straightforward extension of *MSstats*, and is therefore applicable to the full breadth of experiment types supported by *MSstats*. Although demonstrated here on DDA, it is also applicable to DIA, SRM and PRM acquisitions. Additionally, the approach can handle experiments that use different strategies for modified versus unmodified peptides, e.g. using label-free methods for unmodified peptides and TMT labeling for modified peptides, or vice versa.

The proposed approach assumes that all the peptides are correctly mapped to the underlying proteins and PTM sites, and that the features are informative of the protein and PTM abundances. However, multiple modification sites per peptide can confound the abundance of each PTM site. Changes in the unmodified peptide (as opposed to the unmodified protein) can also confound changes in PTM abundance. One potential solution is to quantify the abundance of peptides with one modification and use this to adjust the peptide with multiple sites to remove the confounding. However, this method would likely be challenged by scarsity of modified peptide features containing both a single and multiple modification sites. As the result, peptides with multiple modifications are currently beyond the *MSstatsPTM* scope.

Overall, *MSstatsPTM* balances accuracy and practicality, and enables the analysis of complex experiments in high throughput. Future work is to carry out the inference and testing for not only the relative change of PTM abundance, but also the fraction of the protein that is modified at the particular site (site occupancy, or stoichiometry), and attempt to remove the confounding of individual PTMs in peptides with multiple modifications.

## Acknowledgments

This work was supported in part by NSF DBI-1759736 and Chan-Zuckerberg Essential Open Source Software Award to O.V.

## Abbreviations

MS: mass spectrometry
PTM: Post-translational modifications
TMT: Tandem mass tag
FDR: false discovery rate
PPV: positive predictive value
ANOVA: Analysis of bariance
DDA: Data-dependent acquisitions
DIA: Data-independent acquisition
SRM: Selected reaction monitoring
PRM: Parallel reaction monitoring
KGG: Ub-remnant diglycyl-lysine
GSDMD: Gasdermin D

