## Supplementary for "MSstatsPTM: Statistical relative quantification of post-translational modifications in bottom-up mass spectrometry-based proteomics"

### MSstatsPTM statistical relative quantification of post-translational modifications in global proteomics experiments
### **Supplementary Information**

Devon Kohler[1], Tsung-Heng Tsai[2], Erik Verschueren[4], Ting Huang[1], Trent Hinkle[3], Lilian Phu[3], Meena Choi*[3], and Olga Vitek*[1]

[1]Khoury College of Computer Science, Northeastern University, Boston, MA, USA
[2]Department of Mathematical Sciences, Kent State University, Kent, OH, USA
[3]MPL, Genentech, South San Francisco, CA, USA
[4]ULUA BV, Arendstraat 29, 2018 Antwerp, Belgium
*Corresponding Authors

### 1 Details of the proposed approach

| | | Model | Estimated Log-fold change | Theoretical variance | Estimated variance | Degrees of freedom |
|---|---|---|---|---|---|---|
| **Label-free** ($y_{ij}$ is $\log_2$ intensity in Condition $i$ and BioReplicate $j$) | **Group comparison** | $y_{ij} = \mu_i + \varepsilon_{ij}$ <br> $\sum_{i=1}^{I} \mu_i = 0$ <br> $\varepsilon_{ij} \sim iid\ N(0, \sigma^2)$ | $\bar{Y}_{i.} - \bar{Y}_{i'.}$ | $\frac{2\sigma^2}{I}$ | $\frac{2J \sum_{i=1}^{I} (y_{ij} - \bar{y}_{i.})^2}{J(IJ - I)}$ | $IJ - I$ |
| | **Time course** | $y_{ij} = \mu_i + BioReplicate_j + \varepsilon_{ij}$ <br> $\sum_{i=1}^{I} \mu_i = 0$ <br> $BioReplicate_j \sim iid\ N(0, \sigma_J^2)$ <br> $\varepsilon_{ij} \sim iid\ N(0, \sigma^2)$ | $\bar{Y}_{i.} - \bar{Y}_{i'.}$ | $\frac{2\sigma^2}{I}$ | $\frac{2 \sum_{i=1}^{I} \sum_{i=1}^{J} (y_{ij} - \bar{y}_{i.} - \bar{y}_{j.} + \bar{y}_{..})^2}{J(I-1)(J-1)}$ | $(I-1)(J-1)$ |
| **TMT** ($y_{mij}$ is $\log_2$ intensity in Condition $i$ and BioReplicate $j$ from Mixture $m$) | **Group comparison** | $y_{mij} = \mu_i + Mixture_m + \varepsilon_{mij}$ <br> $\sum_{i=1}^{I} \mu_i = 0$ <br> $Mixture_m \sim iid\ N(0, \sigma_M^2)$ <br> $\varepsilon_{mij} \sim iid\ N(0, \sigma^2)$ | $\bar{Y}_{.i.} - \bar{Y}_{.i'.}$ | $\frac{2\sigma^2}{MJ}$ | $\frac{2J \sum_{i=1}^{I} \sum_{m=1}^{M} (y_{mij} - \bar{y}_{m..} - \bar{y}_{.i.} + \bar{y}_{...})^2}{MJ(MIJ - I - M + 1)}$ | $MIJ - I - M + 1$ |
| | **Time course** | $y_{mij} = \mu_i + BioReplicate_{jm} + \varepsilon_{mij}$ <br> $\sum_{i=1}^{I} \mu_i = 0$ <br> $BioReplicate_{jm} \sim iid\ N(0, \sigma_J^2)$ <br> $\varepsilon_{mij} \sim iid\ N(0, \sigma^2)$ | $\bar{Y}_{.i.} - \bar{Y}_{.i'.}$ | $\frac{2\sigma^2}{MJ}$ | $\frac{2J \sum_{j=1}^{I} \sum_{m=1}^{M} (y_{mij} - \bar{y}_{mj.} - \bar{y}_{.i.} + \bar{y}_{...})^2}{MJ(I-1)(MJ-1)}$ | $(I-1)(MJ-1)$ |

Figure S1: Different models that are fit depending on the experimental design (group comparison and time course) and quantification workflow (label-free versus TMT). The table shows the true values of the standard errors, along with their estimates and the associated degrees of freedom. The same formulas holds when comparing changes in PTM, or hcnges in the unmodified portion of the protein. When combining the two comparisons for an adjustment, the variance must be multiplied by 2.

### 2 Details on the generation of simulated datasets

#### 2.1 Dataset 1 : Computer simulation 1 - Label-free

In the first simulation an experiment with many features per PTM and unmodified protein was created. Additionally this simulation contained no missing data.

- Mean of log-intensity: 25
- Standard deviations of log-intensities for modified and unmodified peptides: 0.2, 0.3
- Difference in PTM abundance between conditions: 0, 1., 2., 3.
- Difference in protein abundance between conditions: 0, 1., 2., 3.
- Number of replicates: 2, 3, 5, 10
- Number of conditions: 2, 3, 4
- Number of realizations: 1000
- Number of features per PTM: 10
- Number of features per unmodified protein: 10
- Missing data: no missing value

#### 2.2 Dataset 2 : Computer simulation 2 - Label-free missing values and low features

In the second simulation we introduced limited feature observations per PTM as well as masking a portion of the observation to simulate missing values.

- Mean of log-intensity: 25
- Standard deviations of log-intensities for modified and unmodified peptides: 0.2, 0.3
- Difference in PTM abundance between conditions: 0, 1., 2., 3.
- Difference in protein abundance between conditions: 0, 1., 2., 3.
- Number of replicates: 2, 3, 5, 10
- Number of conditions: 2, 3, 4
- Number of realizations: 1000
- Number of features per PTM: 2
- Number of features per unmodified protein: 10
- Missing data: 20% of the observations for PTMs and Proteins were masked with NA at random

### 3 Additional evaluation results

#### 3.1 Simulated Datasets 1 and 2

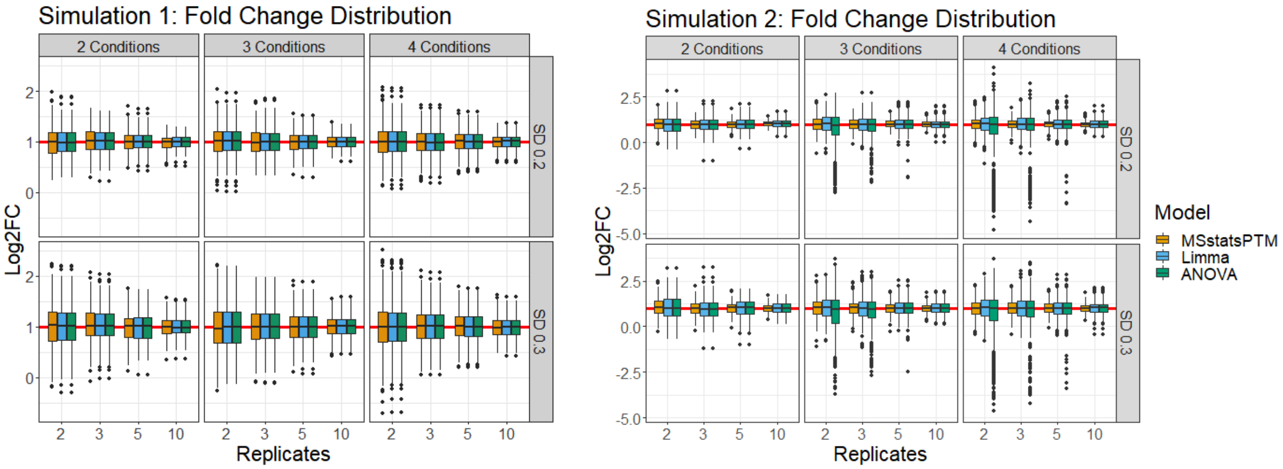

(a) (b)

Figure S2: Simulated Datasets 1 and 2 : fold change distributions. All models are shown after adjusting for changes in unmodified protein abundance. (a) In Simulated Dataset 1 all considered methods correctly estimated the fold change between conditions, with a median fold change estimation of 1. The distributions around the median were also consistent across all methods. (b) In Simulated Dataset 2 all methods correctly estimated the fold change with a median log change of 1. *MSstatsPTM* in this simulation had a tighter distribution around the median. Both *Limma* and *ANOVA* showed a wider range around the fold change.

#### 3.2 Dataset 3 : SpikeIn benchmark - Ubiquitination - Label-free

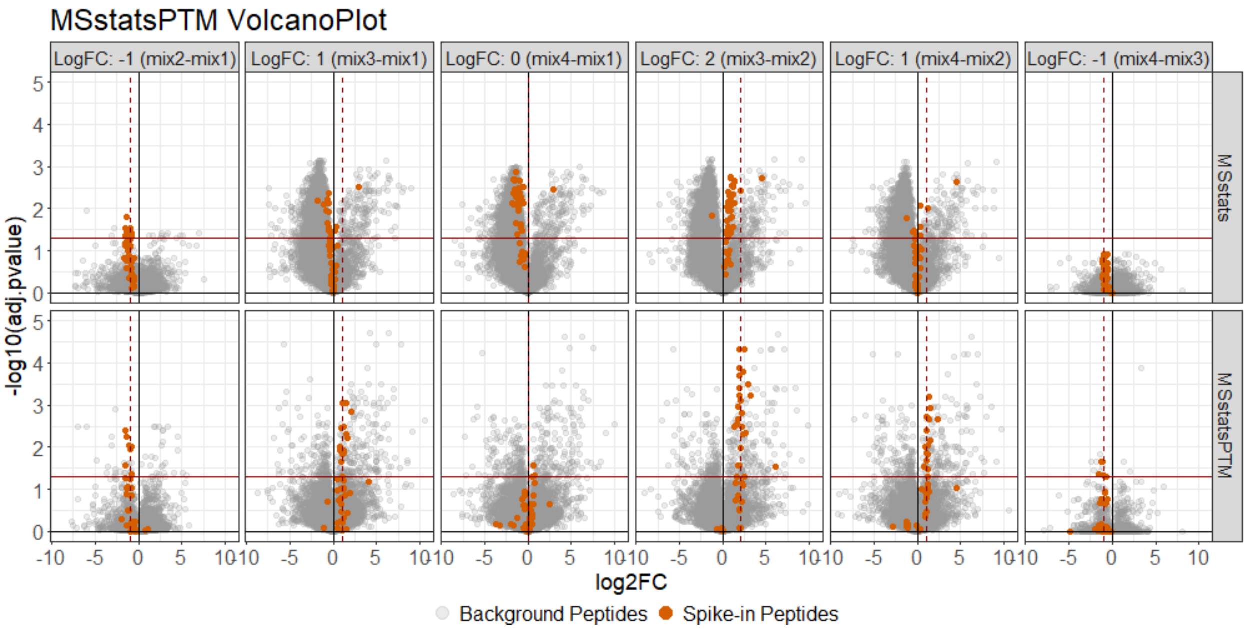

Figure S3: Dataset 3 : SpikeIn benchmark - Ubiquitination - Label-free. The modeling results of the *MSstatsPTM* both before and after adjustment. The unadjusted model is the basic version of *MSstats* before adjustment. The spike-in peptides are colored red and the background peptides are colored grey. All grey peptides are expected to not be detected as differentially abundant. The spike-in peptides (colored red) did not follow the expected log fold change before adjustment. After adjusting for changes in overall protein abundance the spike-in peptides were more in line with expectation. Additionally the background grey colored peptides showed many false positives before adjustment. After adjustment these false positives were decreased considerably.

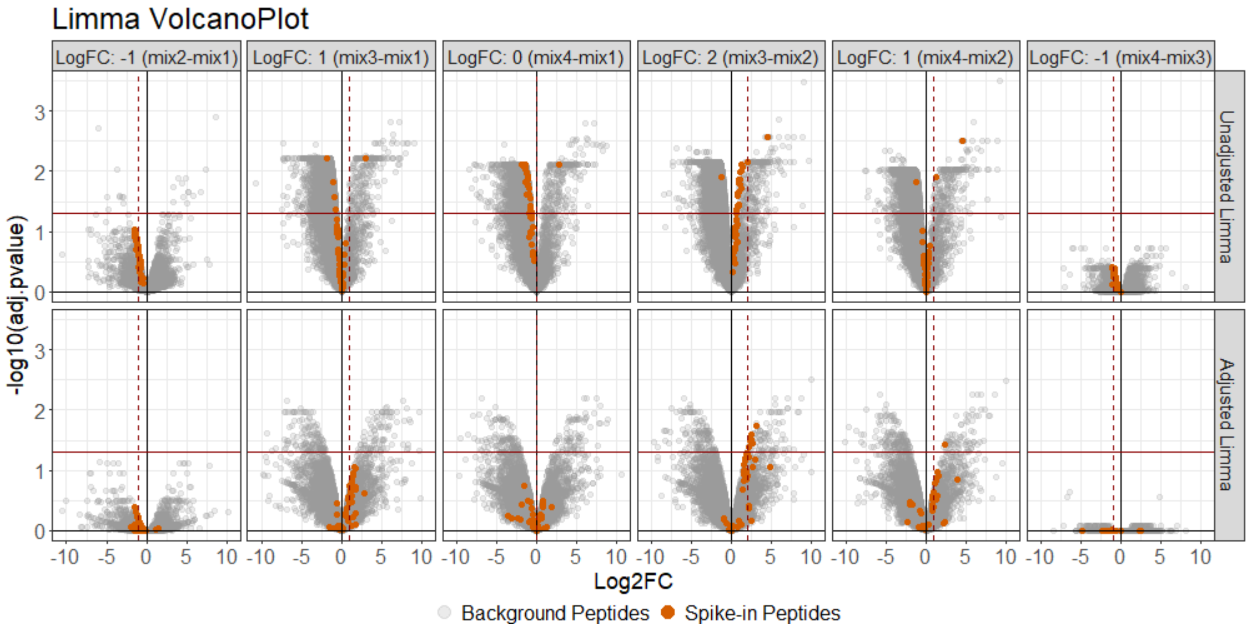

Figure S4: Dataset 3 : SpikeIn benchmark - Ubiquitination - Label-free. When modeling the experiment with the *Limma* method, the spike-in peptides again follow the expected log fold change better after adjusting for changes in protein level. However, while the fold change was more accurate, the majority of spike-in peptides were not detected as differentially abundant. There were more false positive differentially abundant PTM before than after adjustment.

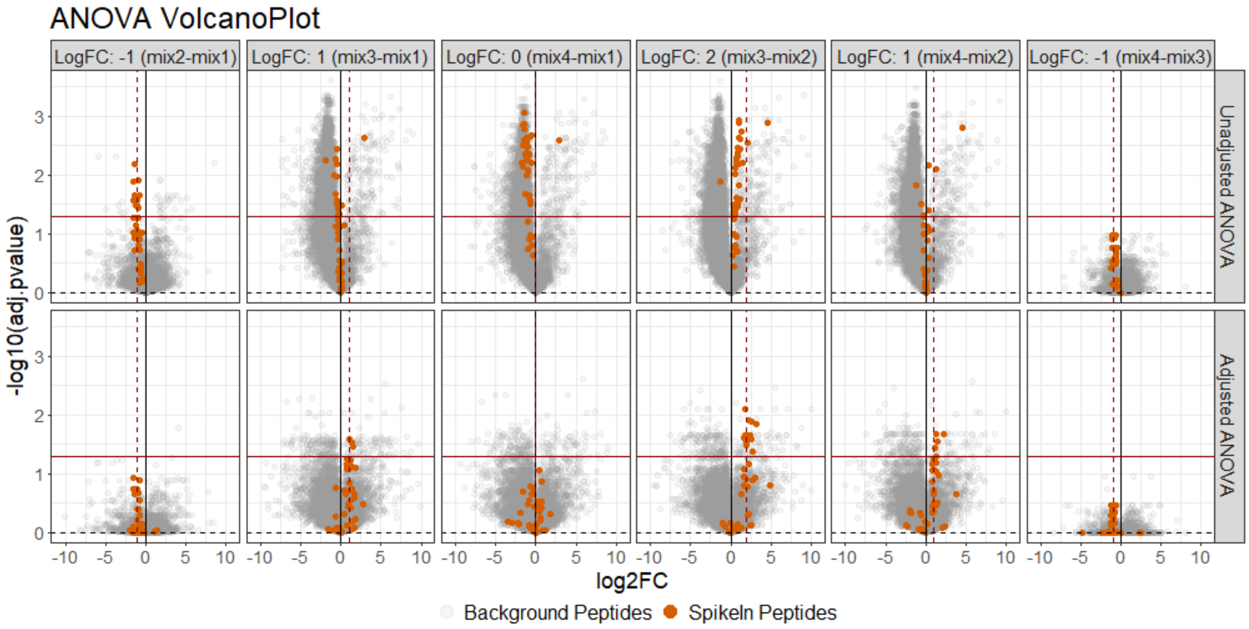

Figure S5: Dataset 3 : SpikeIn benchmark - Ubiquitination - Label-free. Using $ANOVA$ the fold change of the spike-in peptides was much closer to expectation after adjusting for global protein abundance. The log FC estimation was the same as *Limma*, however the p-values were different. In this particular case we detect a few more true positives using $ANOVA$ compared to *Limma*.

##### 3.3 Dataset 4 : Human - Ubiquitination - 1mix-TMT

The experiment had a simple group comparison design, shown in Table S1.

| Condition | BioReplicate | Channel |
|---|---|---|
| Dox1hr | Dox1hr_1 | 127C |
| Dox2hr | Dox2hr_1 | 128N |
| Dox2hr | Dox2hr_2 | 130C |
| Dox4hr | Dox4hr_1 | 128C |
| Dox4hr | Dox4hr_2 | 131C |
| Dox6hr | Dox6hr_1 | 129N |
| Dox6hr | Dox6hr_2 | 131N |
| NoDox0hr | NoDox0hr_1 | 126C |
| NoDox0hr | NoDox0hr_2 | 129C |
| NoDox6hr | NoDox6hr_1 | 127N |
| NoDox6hr | NoDox6hr_2 | 130N |

Table S1: The experimental design of Dataset 4

The following model was fit separately for ubiquitinated peptides and for unmodified protein

$$Y_{mij} = \mu_i + \epsilon_{mij},\ \sum_{i=1}^{I} \mu_i = 0,\ \epsilon_{mij}\ \sim N(0, \sigma^2)$$

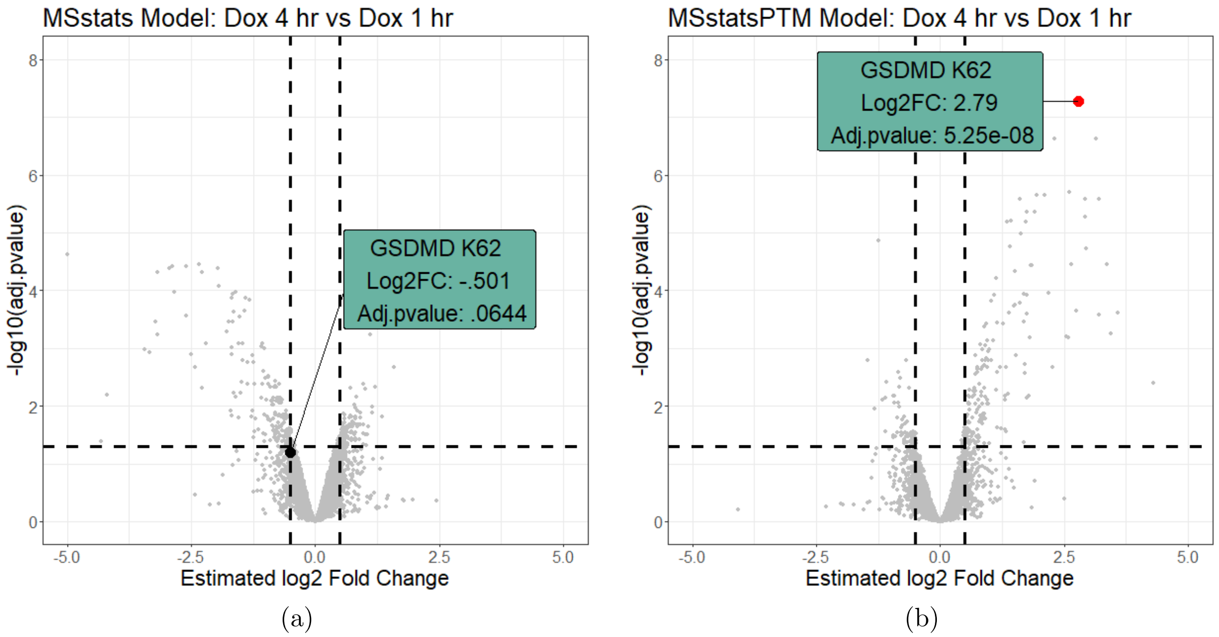

Figure S6: Dataset 4 : Human - Ubiquitination - 1mix-TMT. Volcano plots of Dox4hr vs Dox1hr both before and after protein adjustment with $MSstatsPTM$. The $GSDMD\_HUMAN|P57764\_K62$ modification is highlighted. (a) Before adjustment the modification had a small fold change and was not detected as differentially abundant. (b) After adjustment the fold change was much larger, and the modification was detected as differentially abundant. In this case $MSstatsPTM$ allowed us to identify a differential modified peptide that could have otherwise been missed.

#### 3.4 Dataset 5 : Mouse - Phosphorylation - 2mix-TMT

The experiment had a group comparison design, and the data were acquired in two mixtures, as shown in Table S2.

| | Mixture 1 | | Mixture 2 | | Condition |
|---|---|---|---|---|---|
| Uninfected | 128C | | 128C | 131C | |
| Early (1 Hour) | 126C | 129C | 126C | 129C | WT |
| Late (3 Hour) | 127C | 130C | 127C | 130C | |
| Uninfected | 129N | 131C | 129N | | |
| Early (1 Hour) | 127N | 130N | 127N | 130N | KO |
| Late (3 Hour) | 128N | 131N | 128N | 131N | |

Table S2: The experimental design of Dataset 5

The following model was fit separately for phosphorylated peptides and for unmodified protein

$$Y_{mij} = \mu_i + Mixture_m + \epsilon_{mij},\ Mixture_m \ \sim N(0, \sigma_M^2),\ \sum_{i=1}^{I} \mu_i = 0,\ \epsilon_{mij} \ \sim N(0, \sigma^2)$$

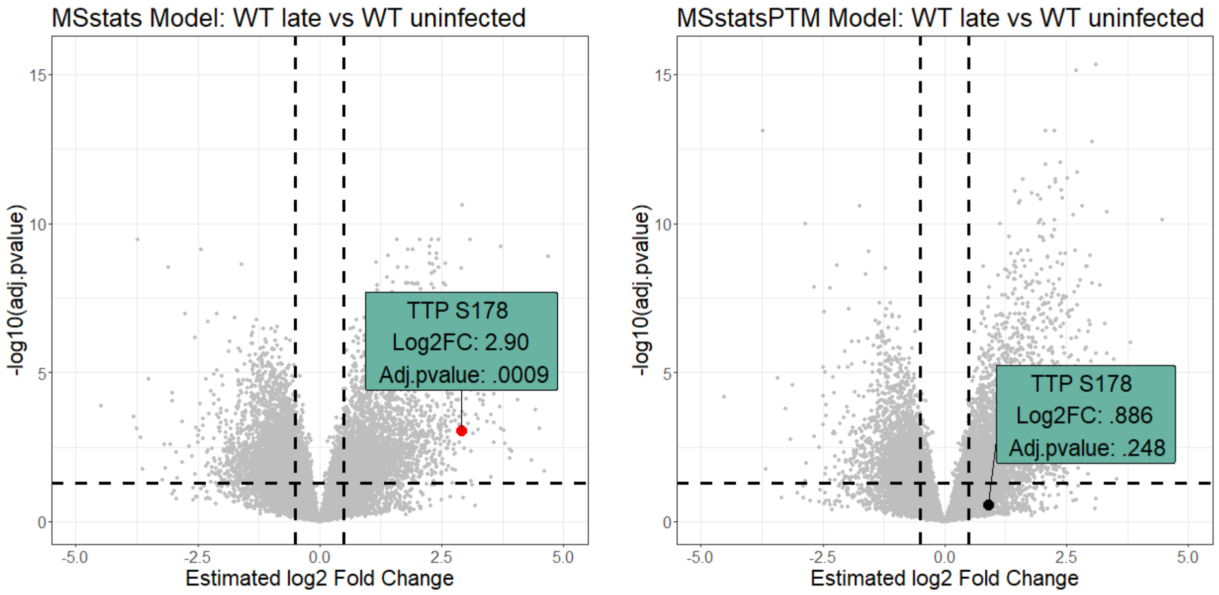

(a) (b)

Figure S7: Dataset 5 : Mouse - Phosphorylation - 2mix-TMT. Volcano plots of WT_Late vs WT_Uninfect both before and after protein adjustment with $MSstatsPTM$. The $TTP\_MOUSE|P22893\_S178$ modification is highlighted. (a) Before adjustment the modification had a large fold change and a small p-value. (b) After adjustment the fold change was much smaller and the modification was not detected as differentially abundant.
